# More than just a passive brick in the wall: the nucleosome facilitates DNA polymerase β activity in linker DNA and its PARP-dependent regulation in the BER sub-pathway choice

**DOI:** 10.64898/2026.02.12.705504

**Authors:** Danil M. Shtanov, Tatyana A. Kurgina, Mikhail M. Kutuzov, Konstantin N. Naumenko, Alexander A. Ukraintsev, Nina A. Moor, Olga I. Lavrik

**Affiliations:** Knorre Institute of Chemical Biology and Fundamental Medicine, SB RAS, Novosibirsk, Russia

**Keywords:** DNA repair, BER, nucleosome, PARP1, PARP2, H1, chromatin, SP-BER, LP-BER

## Abstract

DNA polymerase β (Polβ) is a central player of base excision repair (BER), performing gap-filling synthesis on damaged DNA. While nucleosome core particles (NCPs) are known to impede activity of BER enzymes, the regulation of this process in linker DNA adjacent to nucleosomes remains unclear. Here we demonstrate an unexpected stimulation of Polβ-catalyzed strand-displacement synthesis in linker DNA by the adjacent NCP. Notably, the nucleosome context reinforces the regulatory modulation of Polβ activity by PARP1/PARP2 and FEN1. While linker histone H1 restricts strand-displacement synthesis at the nucleosome entry/exit site, PARP1 and PARP2 modulate Polβ function through competitive binding to DNA gaps or nicks and via poly(ADP-ribosyl)ation (PARylation). At the same time, PARPs binding differentially regulates BER sub-pathway choice, and PARylation alleviates H1-mediated inhibition. These findings reveal a multi-layered regulatory system wherein the nucleosome acts as a dynamic platform coordinating Polβ activity and its interplay with chromatin-associated factors, influencing the balance between short- and long-patch BER. The research advances understanding of chromatin-mediated control of BER DNA repair synthesis and the functional specialization of PARP1 and PARP2 in maintaining genome stability.

**Highlights:** The nucleosome core particle acts not only as a barrier but also as a stimulator of Polβ-mediated DNA repair synthesis in adjacent linker DNA.

The nucleosome acts as a platform that strengthens the regulation of repair synthesis in linker DNA by chromatin-associated factors (PARP1, PARP2, H1).

PARP1 suppresses overall Polβ synthesis, while PARP2 specifically inhibits strand-displacement, thereby gating the choice between short- and long-patch BER sub-pathways.

## Introduction

DNA polymerase β (Polβ) is the primary DNA polymerase responsible for gap-filling synthesis during the base excision repair (BER)^1,2^. This enzyme processes the central 1-nucleotide (nt) gap intermediate through two distinct activities: its polymerase domain catalyzes nucleotide insertion, while its N-terminal lyase (8-kDa) domain excises the 5’-deoxyribose phosphate (5’-dRP) moiety, enabling the short-patch (SP) BER sub-pathway. Polβ can also perform limited strand-displacement synthesis, a prerequisite for the alternative long-patch (LP) BER sub-pathway where 2-11 nucleotides are incorporated^3^. A defining biochemical feature of Polβ is its distributive mode of synthesis. Unlike processive replicative polymerases, Polβ does not remain tightly bound to its DNA template but instead dissociates after incorporating a short patch of 2-3 nucleotides^4^. However, this distributive kinetics also means that the overall efficiency of DNA synthesis is highly dependent on the enzyme’s rebinding rate. The resulting flap is cleaved by flap endonuclease 1 (FEN1), which stimulates Polβ activity in strand-displacement synthesis, in accordance with the "passing the baton" mechanism^5–7^. Notably, the fidelity of Polβ decreases substantially during strand-displacement synthesis compared to single-nucleotide gap filling, highlighting the critical importance of regulating switch between the two mechanisms^8^. Overexpression of Polβ, as observed in certain malignancies, is thought to promote a shift towards the long-patch BER (LP-BER) sub-pathway and, under specific conditions, may even facilitate its aberrant recruitment into the nucleotide excision repair (NER) pathway. Such misregulation ultimately contributes to increased mutation rates and genomic instability^9,10^.

The progression and efficiency of BER are tightly regulated by various factors, prominently including poly(ADP-ribose) polymerases 1 and 2 (PARP1 and PARP2). These enzymes function as molecular sensors of DNA strand breaks, binding to damaged DNA and catalyzing the synthesis of poly(ADP-ribose) (PAR) chains^11^. PARylation serves as a signal for the recruitment of DNA repair factors but also modulates enzyme activity; for instance, automodification of PARP1 and PARP2 promotes their dissociation from DNA, thereby facilitating access for repair enzymes like Polβ^12–15^. PARP1 and PARP2 exhibit distinct affinities for different BER intermediates: PARP1 binds effectively to undamaged DNA, AP-sites, gaps and nicks, whereas PARP2 shows a marked preference for gapped and nicked DNAs^12,13,16,17^. Consequently, PARP1 influences all stages of BER^12,18^, while PARP2 acts as a key regulator of final steps, including the handoff between Polβ and DNA ligase IIIα (LigIIIα)^14^.

A critical layer of DNA repair regulation is added by the packaging of genomic DNA into chromatin. The nucleosome core particle (NCP), the fundamental repeating unit of chromatin, presents a significant structural barrier to DNA repair machinery^19^. Extensive biochemical and recent structural studies, comprehensively detailed in an excellent recent review^20^, have demonstrated that the activities of DNA glycosylases, apurinic/apyrimidinic endonuclease 1 (APE1), Polβ, and LigIIIα are substantially suppressed on nucleosomal DNA compared to naked DNA^14,21–24^. The translational and rotational positioning of a DNA lesion within the nucleosome profoundly influences its accessibility to BER enzymes^25–29^. For example, the activity of Polβ in gap-filling synthesis within the nucleosome follows a strong position-dependent gradient, being most efficient near the entry/exit sites^25,30^ or linker DNA^26^. Cryo-EM structures reveal that the reduced activity of Polβ in processing low-accessible lesions stems from the necessity for enzyme to engage in extensive "global DNA sculpting," displacing ∼35 base pairs of DNA from the histone octamer to induce a ∼90° DNA bend^30^. Similarly, ligation of the final nick by the XRCC1-LigIIIα complex is efficient only near the nucleosome periphery^23^. This constrained repair capacity within chromatin is biologically significant, as it correlates with decreased DNA repair efficacy and mutation rates in nucleosome-dense genomic regions *in vivo*^31–33^.

To overcome the chromatin barrier, cells employ active chromatin remodeling strategies^34,35^. PARP1 and PARP2, in complex with the histone PARylation factor 1 (HPF1), play a pivotal role in this process by catalyzing PARylation of core histones, primarily on serine residues^36–38^. This HPF1-dependent histone PARylation triggers local chromatin relaxation facilitating the access of repair factors to lesions^39^. While histone PARylation may directly relax nucleosome structure to facilitate Polβ activity on nucleosome substrates *in vitro*^40^, its principal function *in vivo* is likely the targeted recruitment of chromatin remodelers^41–44^. ADP-ribosylation of histones in the vicinity of a nucleosomal DNA lesion, which is most efficiently catalyzed by PARP2^16,17,45^, might serve as a signal to recruit ALC1. This remodeler would then catalyze nucleosome sliding away from the damage site, effectively repositioning the lesion into the more accessible linker DNA, where it becomes fully exposed to the repair machinery^44,46–48^.

Whereas the mechanisms of BER within the nucleosome core have been increasingly elucidated, the repair landscape in linker DNA regions – the stretches connecting nucleosome cores – remains less characterized. The nucleosome itself imposes a strong bias for the SP-BER sub-pathway, effectively limiting DNA repair synthesis within the nucleosome core to single-nucleotide insertion; this phenomenon is less pronounced in linker DNA^49^.

It was previously established that Polβ-catalyzed strand-displacement synthesis in linker DNA produced a repair patch whose size is spatially constrained by the proximity of the nucleosome^49^. However, these investigations, performed in nuclear extracts, do not fully delineate the specific interplay between Polβ and its regulators like PARPs in the context of linker DNA. Furthermore, the entry/exit site of the nucleosome is associated with the linker histone H1, which promotes higher-order chromatin compaction and can further hinder the access of repair enzymes to DNA lesions^22,50^. Recent findings indicate that the eviction of H1 from chromatin facilitates DNA repair, suggesting its dynamics is a key regulatory point^22,51^.

In this study, we investigate the activity of Polβ on linker DNA and its functional interplay with the linker histone H1 and the PARP1/PARP2 system, including the HPF1 cofactor. We specifically focus on how an adjacent nucleosome core particle and its associated proteins regulate the balance between Polβ-catalyzed gap-filling and strand-displacement synthesis, a critical point in the BER sub-pathway choice.

## Results

### Stimulation of Polβ activity in Strand-Displacement Synthesis by an Adjacent Nucleosome Core Particle

Previous studies have consistently demonstrated that Polβ activity is substantially constrained within the nucleosome core, while its behavior on linker DNA adjacent to a nucleosome core particle (NCP) remains less explored. To investigate this aspect, we generated a nucleosome substrate with 50 bp and 30 bp linker arms, containing a 1-nt gap located 26 bp from the 5’ end of the longer linker and 23 bp from the nucleosome boundary (gap27-NCP227) (Fig. 1a). The nucleosome was reconstituted using the Widom 603 positioning sequence, which ensures a well-defined translational and rotational setting of the DNA. Along with the corresponding full-length naked DNA (gap27-DNA227), we prepared a 50 bp duplex (gap27-DNA50) equivalent in both the sequence and the gap position to the linker arm (Fig. 1a). The latter substrate provides the same amount of Polβ-accessible DNA as gap27-NCP227, in which ∼170 bp is wrapped around the histone octamer, and thus controls for a possible sequestration of the enzyme on the extra free DNA present in gap27-DNA227.

**Fig. 1.**
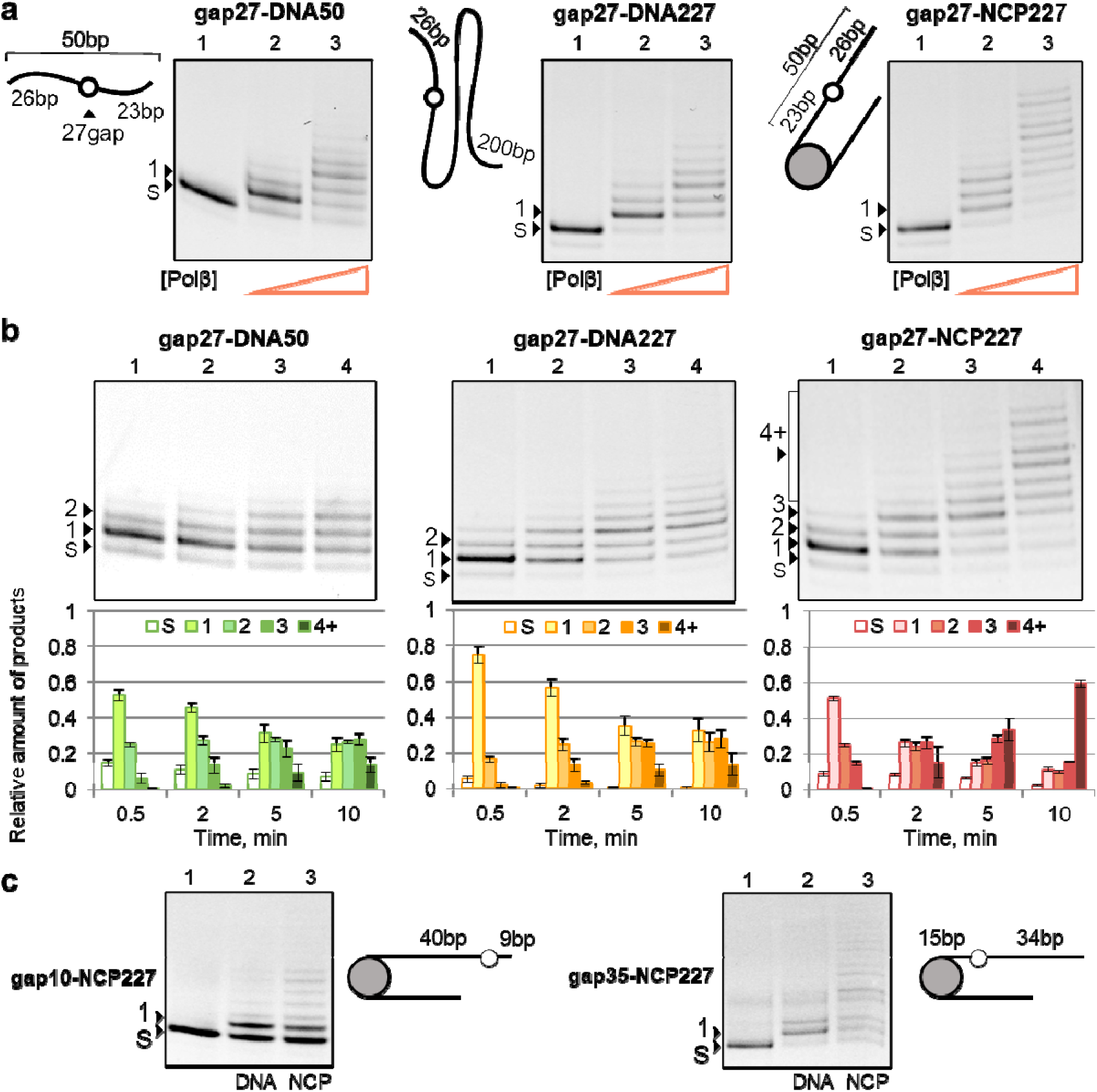
The adjacent NCP stimulates Polβ-catalyzed strand-displacement synthesis in linker DNA. (**a**) Schematic structures of the substrates and electrophoregrams of the strand-displacement synthesis products. Left to right: gap27-DNA50, a 50 bp duplex corresponding to the long linker arm; gap27-DNA227, the full-length naked DNA; gap27-NCP227, the nucleosome assembled on the same 227 bp DNA; the position of 1-nt gap is the same in all three substrates. Each substrate (50 nM) was incubated with all four dNTPs (100 μM each) for 10 min without Polβ (lane 1), with 6.5 nM Polβ (lane 2) or 25 nM Polβ (lane 3). (**b**) Time course of strand-displacement synthesis on gap27-DNA50, gap27-DNA227 and gap27-NCP227. Substrates (50 nM) were incubated with 20 nM Polβ and four dNTPs (100 μM each) for 0.5, 2, 5 and 10 min. Upper panels – representative electrophoregrams; lower panels – relative amounts of the substrate (S) and products of 1, 2, 3 or 4 and more (4+) nucleotides insertion, quantified from independent experiments (mean ± SD, n = 3). (**c**) Effect of the gap position within the linker. Two nucleosome substrates bearing the 1-nt gap farther from (gap10-NCP227, 40 bp from the boundary of NCP) or closer to (gap35-NCP227, 15 bp from the boundary of NCP) the nucleosome were examined. Each substrate (50 nM) was incubated with four dNTPs (100 μM each) and 25 nM Polβ for 10 min: lane 1 – substrate without Polβ; lane 2/3 – naked DNA/NCP with Polβ. The positions of the substrate (S) and extension products (n) in denaturing 15% PAG are indicated to the left of the gel images.

Unexpectedly, we found that the efficiency of Polβ-catalyzed strand-displacement synthesis was higher on gap27-NCP227 substrate compared to the respective naked DNA (gap27-DNA227) and to gap27-DNA50 as well. Under identical reaction conditions, Polβ synthesized markedly longer extension products on gap27-NCP227 than on either naked DNA substrate, with DNA extension products generated from gap27-DNA50 and gap27-DNA227 being very similar in the length and relative yields (Fig. 1a). Kinetic experiments also show more efficient strand-displacement synthesis on gap27-NCP227 substrate at each time point (Fig. 1b); after 10-min incubation, the relative yield of fraction 4+ corresponding to the insertion of 4 and more nucleotides (nts) was the highest on gap27-NCP227, while the main products of gap27-DNA50 and gap27-DNA227 conversion corresponded to the insertion of 1–3 nts.

In contrast to the strand-displacement synthesis, the single-nucleotide gap-filling reaction catalyzed by Polβ was observed to proceed on all three substrates at closely similar rates (Supplementary Fig. 1). The insertion of a single nucleotide under these conditions was additionally confirmed by using ddTTP as a chain-terminating substrate. Thus, modulation of Polβ activity in strand-displacement synthesis by the adjacent NCP is related to the DNA flap formation rather than to insertion of the first nucleotide.

To examine whether the stimulatory action of NCP depends on the location of the lesion within the linker, we constructed NCP structures carrying the gap farther (gap10-NCP227) and closer (gap35-NCP227) to the nucleosome boundary. In both cases the strand-displacement synthesis remained more efficient on the nucleosome substrate than on the respective naked DNA (Fig. 1c), indicating that the observed stimulation by the NCP is a general property of linker DNA repair.

A potential factor that may influence the efficiency of strand-displacement synthesis in linker DNA is interaction between the displaced single-stranded DNA flap (5’-flap) and the NCP. It is plaus ble that the flexible N-terminal tails of core histones could interact with the displaced flap, thereby facilitating destabilization and unwinding of DNA helix and enhancing the synthesis. Such an interaction would not require the nucleosome to be covalently continuous with the damaged DNA. To test this, we supplied the reaction mixture with an undamaged NCP227 as a trans-acting factor, at a concentration equimolar to the gap-DNA/gap-NCP substrate. The addition of NCP227 stimulated strand-displacement synthesis on the naked gap27-DNA227: the fraction of products with 4–7 inserted nts increased approximately four-fold at the expense of the shorter products, and the insertion of 8 and more nts (fraction 8+) became detectable (Fig. 2a). However, the yield of the longest products was lower vs that on gap27-NCP227 itself (compare samples 3 and 4). Consistently, addition of NCP227 to gap27-NCP227 produced no stimulatory action (compare samples 4 and 5).

**Fig. 2.**
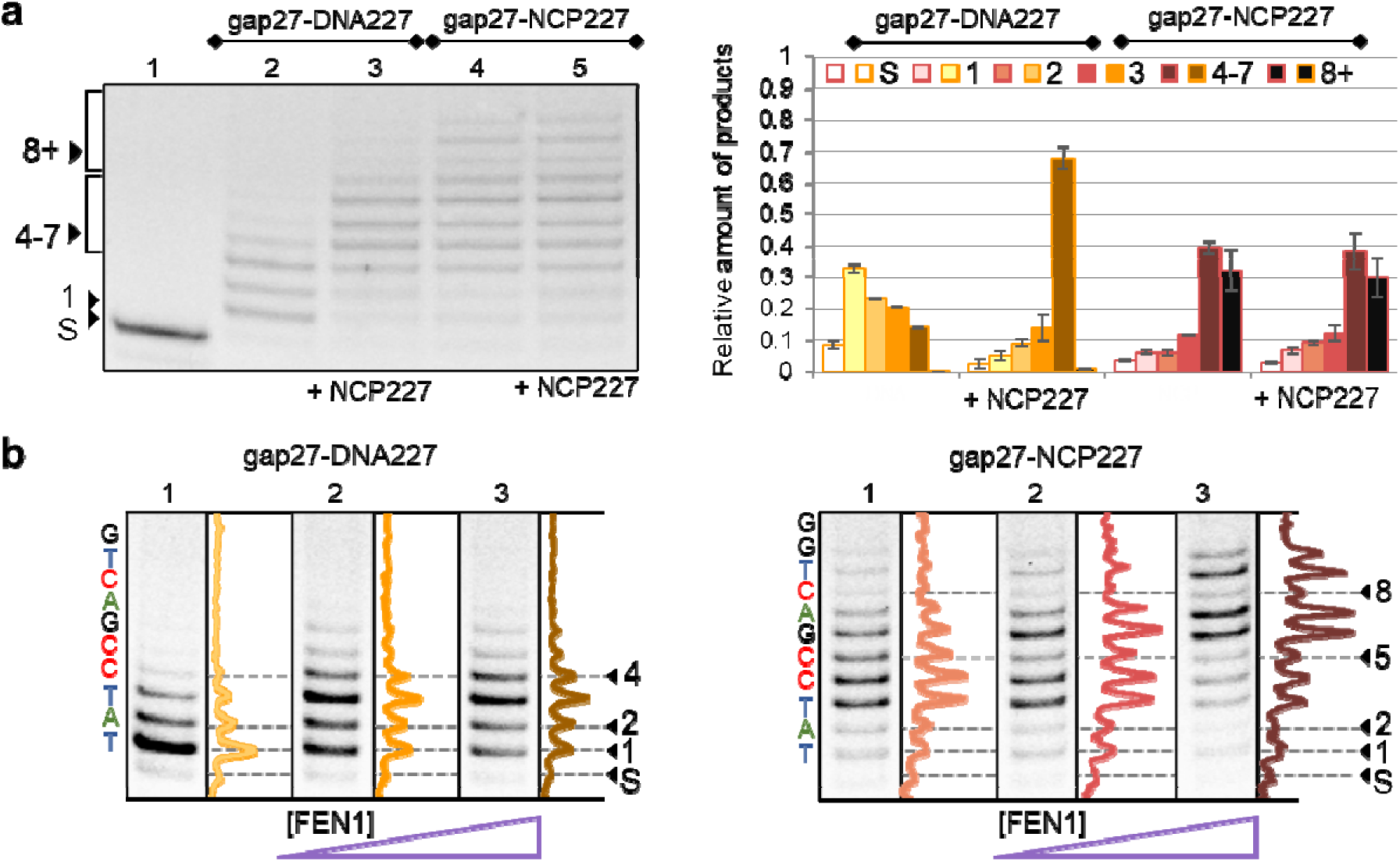
Strand-displacement synthesis in the linker region of gap-NCP is modulated in a specific manner by the nucleosome and FEN1. (**a**) Impact of the non-gapped NCP. 50 nM gap27-DNA227 (lanes 2, 3) or gap27-NCP227 (lanes 4, 5) was incubated with 20 nM Polβ and four dNTPs (100 μM each) for 10 min in the absence (lanes 2, 4) or presence (lanes 3, 5) of 50 nM non-gapped NCP227. Relative amounts of the substrate (S) and extension products determined from independent experiments (mean ± SD, n = 3) are shown. (**b**) Influence of FEN1. 50 nM gap27-DNA227 or gap27-NCP227 was incubated with 20 nM Polβ and four dNTPs (100 μM each) for 10 min with increasing concentrations of FEN1 (lane 1 – no FEN1; lane 2/3 with 25/50 nM FEN1 for each substrate). The displaced strand sequence downstream of the gap is shown to the left of each gel; the densitometric profile of each lane is shown to the right. Note the periodic attenuation of the bands corresponding to every third inserted nucleotide (products 2, 5, 8) detected on the nucleosome substrate only.

Two further control experiments argue that stimulation of strand-displacement synthesis requires the assembled particle rather than the mere presence of protein(s) in the reaction mixture. Addition of BSA to either gap27-DNA227 or gap27-NCP227 did not affect the activity of Polβ in DNA synthesis (Supplementary Fig. 2a), ruling out possible nonspecific enzyme stabilization in the presence of trans-acting NCP. Addition of free core histones to gap27-DNA227 at varied concentrations did not stimulate Polβ activity (Supplementary Fig. 2b). Instead, free histones present at excess over Polβ suppressed strand-displacement synthesis, likely due to competition with Polβ for the substrate binding via nonspecific electrostatic interaction with the DNA. Accordingly, our data show that the assembled nucleosome stimulates strand-displacement synthesis even when acting in trans, but they do not resolve whether the histone tails are directly involved – a question that would require experiments with nucleosomes reconstituted from tail-truncated histones.

The displaced 5’-flap is not merely a by-product of strand-displacement synthesis: it is the key intermediate of the LP-BER sub-pathway and a substrate of flap endonuclease 1 (FEN1), which cleaves it off to allow subsequent ligation. Consequently, if the adjacent nucleosome indeed acts on the displaced strand, it should affect not only the extent of synthesis but also its coordination with the flap removal. We therefore analyzed the Polβ polymerase activity in the presence of FEN1. Consistent with previous reports^7^, the addition of FEN1 stimulated overall DNA synthesis by Polβ on both naked gap27-DNA227 and gap27-NCP227 substrate, as evidenced by an increased yield of long extension products (Fig. 2b). This is attributed to FEN1 resolving inhibitory secondary structures in the displaced flap, allowing Polβ to perform more processive synthesis. Strikingly, a particular pattern of products emerged specifically on the nucleosome substrate. The electrophoretic profile revealed a pronounced attenuation of bands corresponding to the incorporation of every third nucleotide (products 2, 5, 8), creating a periodic pattern (Fig. 2b, gap27-NCP227). In contrast, the product distribution detected for the naked DNA was more uniform, without the pronounced periodic intensity pattern (Fig. 2b, gap27-DNA227). This observation suggests that in the nucleosome context, the cleavage of the displaced flap by FEN1 is exceptionally efficient and tightly coupled to the synthetic cycle of Polβ. We hypothesize that upon the formation of a 3-nt flap – a preferred substrate for FEN1 in the nucleosome substrate – the endonuclease immediately cleaves it, effectively "resetting" the substrate and allowing Polβ to rapidly initiate the next synthetic cycle. This highly coordinated cycle results in the observed rhythmic extension pattern, illustrating a processive "passing the baton" mechanism uniquely facilitated by the nucleosome scaffold.

To determine whether this coupling between Polβ and FEN1 manifested in the rhythmic extension pattern requires the cis-acting nucleosome, we explored the impact of FEN1 on Polβ activity in the absence and presence of non-gapped NCP227 (Supplementary Fig. 3). On gap27-DNA227, the trans-acting nucleosome stimulated production of long products but did not induce the periodic attenuation pattern; the addition of NCP227 to gap27-NCP227 attenuated the periodicity driven by FEN1. Thus, the trans-acting nucleosome can stimulate activity of Polβ in the strand-displacement synthesis without affecting its coupling to FEN1, which requires the inherent nucleosome.

### Interaction of Polβ and its Possible Competitors with Gapped DNA and Nucleosome Substrates

The stimulation of strand-displacement synthesis we observed could theoretically be attributed to enhanced affinity of Polβ for the nucleosome substrate, which would accelerate the rebinding of this enzyme characterized by a distributive mechanism of action. Moreover, Polβ is not the only protein that can interact with damaged DNA in the linker region. Linker histone H1 binds to the dyad region between the entry and exit sites of nucleosome-wrapped DNA; PARP1 and PARP2 act as sensors of DNA strand breaks. Thus, a quantitative comparison of their affinities for the same substrates is essential for interpreting their functional interplay with Polβ, as described in subsequent sections. We compared binding affinities of Polβ and its possible competitors (linker histone H1, PARP1 and PARP2) for gap27-DNA227, gap27-NCP227, and nick-NCP227 (the product of 1-nt gap-filling synthesis), using fluorescence titration assay (Supplementary Fig. 4a–d).

As shown by apparent equilibrium dissociation constants of complexes determined as EC₅₀ values (Table 1), the linker histone H1 exhibits the lowest affinity for all the substrates, though its affinity for the NCP is 1.5-fold higher than for the naked DNA (*p* ≤ 0.05), consistent with its specific binding at the nucleosome entry/exit site^52–54^. PARP1 and PARP2 exhibited comparable affinity for gap27-DNA227 and gap27-NCP227. However, the PARP2 affinity for nick-NCP227 was higher than that of either PARP1 or Polβ (*p* ≤ 0.01 and 0.05, respectively), which is consistent with previously published data^13,14^. As expected, Polβ exhibited a higher affinity for the gapped substrates than for the single nucleotide insertion product, nick-NCP (*p* ≤ 0.05). It is noteworthy that the affinity of Polβ for gap27-NCP227 was only 1.5-fold higher than its affinity for gap27-DNA227 – that is the difference was as small as that observed for the gap-filling rate constants (Supplementary Fig. 1d); consequently, this is insufficient to explain the several-fold stimulation of strand-displacement synthesis on the nucleosome substrate.

**Table 1.** EC₅₀ values (nM) for protein complexes with DNA and nucleosomes.

| Substrate | PARP1 | PARP2 | H1 | Polβ |
| --- | --- | --- | --- | --- |
| gap27-DNA227 | 3 ± 0.3 | 3 ± 0.4 | 12 ± 2 | 1.5 ± 0.3 |
| gap27-NCP227 | 3 ± 0.4 | 2 ± 0.3 | 8 ± 1 | 1 ± 0.2 |
| nick-NCP227 | 4 ± 0.6 | 1 ± 0.2 | 8 ± 1 | 2 ± 0.4 |

Statistically significant differences between EC<sub>50</sub> values were evaluated by Student *t*-test; respective *p* values are presented in the text.

Next, we examined Polβ binding to NCP and gap-NCP using mass photometry (MP), which allows quantitative analysis of biomolecular assemblies and their stoichiometries^55^. These data are helpful for correct design and faithful interpretation of experiments performed at different enzyme concentrations. The theoretical mass of Polβ is 39 kDa, and of the nucleosome is 248 kDa (108 kDa histone octamer + 140 kDa DNA). Free Polβ was registered at 69 ± 15 kDa (Fig. 3), consistent with its mass being close to the lower detection limit of mass photometry (∼40 kDa), where accuracy degrades and apparent masses are biased upward^56,57^. However, the observed mass increments for the gap-NCP-associated species correspond exactly to the mass of a single Polβ molecule (Fig. 3b, c). At a 3:1 ratio of Polβ and gap27-NCP227 concentrations, we detected a complex with the mass of 281 ± 15 kDa, indicating a 1:1 ratio of binding (Polβ•gap-NCP227). Complexes of 325, 371 and 414 kDa formed at increased Polβ:gap-NCP227 ratio contained two, three and four Polβ molecules.

**Fig. 3.**
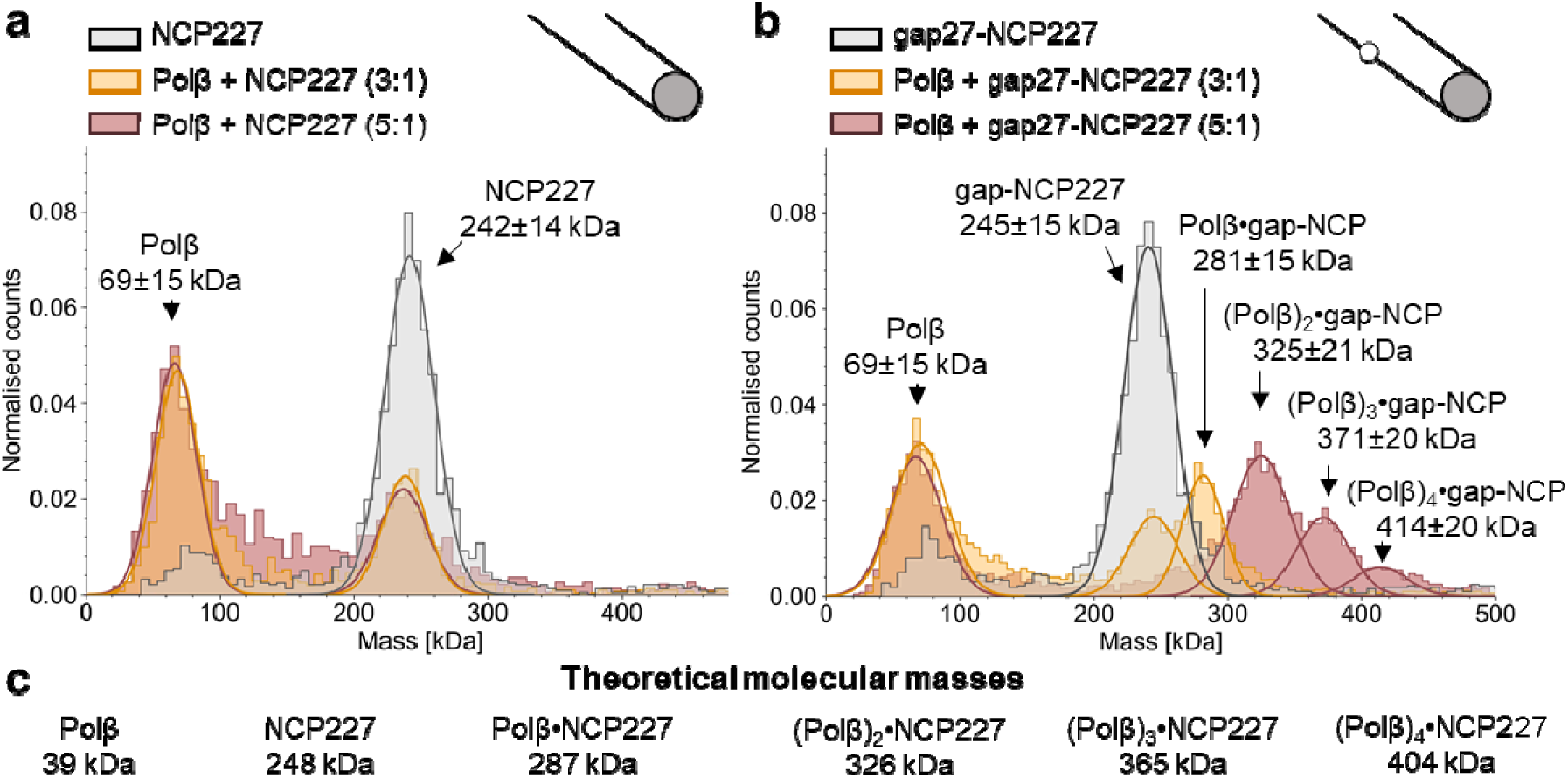
Polβ forms equimolar and higher-order complexes with the gapped nucleosome. (a, b) Mass photometry data: molecular mass distributions of species in samples containing 5 nM NCP227 (**a**) or gap27-NCP227 (**b**), alone or mixed with Polβ at 3:1 and 5:1 protein-to-substrate molar ratios. Gaussian fits and the corresponding mass values (mean ± SD) are shown for the resolved species. The peaks observed for NCP227 in the mixtures with Polβ at any of the ratios correspond to the individual components. Additional species observed for gap27-NCP227 in the mixtures with Polβ correspond to the indicated complexes. (**c**) Theoretical masses of Polβ, NCP227 and their complexes of different stoichiometries for comparison with the experimental values.

Additional binding experiments performed using EMSA revealed the formation of higher-order complexes of Polβ with gap27-DNA227 and gap27-NCP227 (Supplementary Fig. 4e), confirming the data obtained by MP. Importantly, we observed no stable complexes of Polβ with the non-gapped DNA227 and NCP227 substrates in MP and EMSA experiments (Fig. 3a; Supplementary Fig. 4e). These results suggest that i) the stimulation of strand-displacement synthesis by trans-acting NCP is unlikely to result from the direct capture of Polβ by the nucleosome but rather indicates an effect on the DNA itself; ii) the cooperative binding is characteristic of Polβ interaction with its specific (gapped) substrate. In this regard, it should be noted that all DNA synthesis experiments were performed at Polβ concentrations not exceeding the substrate concentration in order to minimize the contribution of higher-order complexes.

### Linker Histone H1 Restricts Strand-Displacement Synthesis by Polβ at the Nucleosome Entry/Exit Site

Linker histone H1 binds to linker DNA and facilitates chromatin compaction, which could potentially inhibit activities of DNA-dependent enzymes near the NCP boundary. We therefore analyzed the influence of H1 on Polβ-catalyzed strand-displacement synthesis, using two substrates bearing gap at different positions. In gap27-NCP substrate, the incorporation of 24 nucleotides corresponds to synthesis within the linker DNA, and the subsequent synthesis proceeds into the nucleosomal DNA at superhelical locations (SHLs); for gap35-NCP227, the boundary corresponds to 16 nts. While previous studies established severe constraints for strand-displacement synthesis within the NCP itself^20^, here we detected the DNA synthesis initiating in the linker region and progressing into the nucleosome (Fig. 4a, lanes 4).

**Fig. 4.**
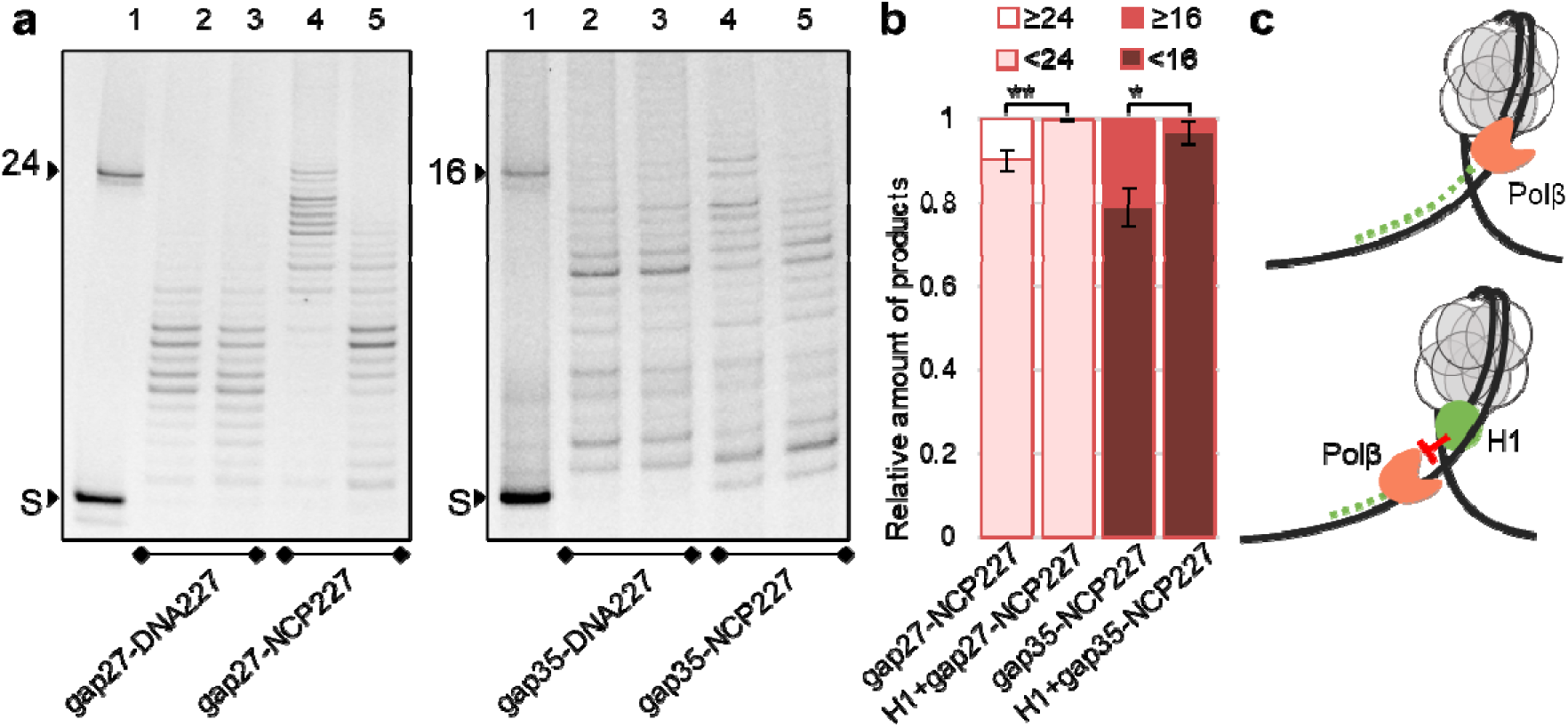
Influence of linker histone H1 on Polβ activity. (**a**) Electrophoregrams show DNA extension via strand-displacement synthesis after incubation of 150 nM gap-DNA/gap-NCP with 100 nM Polβ and four dNTPs (100 μM each), without or with 300 nM H1 for 10 min, for the gap located 23 bp (gap27) or 15 bp (gap35) from the nucleosome boundary. Lane 1 – substrate and marker of DNA extension length; lane 2/3 – gap-DNA without/with H1; lane 4/5 – gap-NCP without/with H1. Positions of the substrates and products in denaturing 15% PAG are indicated to the left of gel images. (**b**) Relative amounts of products divided in two fractions of different lengths as indicated for each nucleosome substrate, without and with H1, quantified from three independent experiments (mean ± SD, n = 3). Statistically significant differences in the yields of longer products in the absence vs presence of H1 are shown (Student’s t-test, *p ≤ 0.05 for gap35-NCP227, **p ≤ 0.01 for gap27-NCP227). (**c**) Scheme shows proposed models of DNA extension by Polβ via strand-displacement synthesis on NCP or its complex with histone H1.

We revealed no impact of H1 on Polβ activity when using naked DNAs (Fig. 4a, lanes 2 and 3). This result can be explained by non-specific binding of H1 to the naked DNA outside the gap region – the site occupied by Polβ due to its higher affinity for the gapped DNA/NCP (Table 1). In contrast, H1 significantly restricted the length of strand-displacement products synthesized on the gap-NCP (Fig. 4a, lanes 5 vs lanes 4), leading to the accumulation of shorter extension products. Notably, the restriction becomes apparent as the synthesis approaches position 24 - the boundary between the linker and the nucleosomal DNA, where H1 specifically binds^52–54^. The same restriction was observed for gap35-NCP, where it appeared at the corresponding position 16 (Fig. 4a, lanes 4, 5); quantification showed that H1 significantly reduced the fraction of products extending beyond the nucleosome boundary on both substrates (Fig. 4b). Thus, while the nucleosome itself promotes more extensive strand-displacement synthesis, the linker histone bound to the same region acts in the opposite direction, limiting the length of the repair patch. Our data indicate that Polβ strand-displacement activity can be regulated not only by the nucleosome core but also by the linker histone H1 (Fig. 4c).

### PARP1 and PARP2 Distinctly Modulate Polβ Activity in Linker DNA Adjacent to the Nucleosome

We further explored the modulation of strand-displacement synthesis induced by PARP1/PARP2 in experiments with a constant Polβ and increasing PARP concentrations. Both PARPs reduced the y eld of strand-displacement products in a concentration-dependent manner; however, as quantitative analysis of the reaction products showed (Fig. 5a, b), they acted at different stages of synthesis. PARP1 suppressed both 1-nt gap-filling and strand-displacement synthesis due to competition with Polβ for the gapped DNA/NCP substrate: the fraction of unreacted substrate progressively accumulated, reaching 0.38 on gap27-DNA227 and 0.60 on gap27-NCP227 at 200 nM PARP1, whereas in its absence the substrate was almost completely consumed. PARP2 was similar to PARP1 in suppression of strand-displacement but did not inhibit the first nucleotide insertion, suggesting that PARP2 competes with Polβ more strongly for binding to the nicked DNA/NCP intermediate. Indeed, a very negligible amount of unreacted substrate was detected at any PARP2 concentration on either substrate; instead, the product of the single nucleotide insertion accumulated, reaching 0.75 on gap27-DNA227 and 0.95 on gap27-NCP227 at 200 nM PARP2, with the strand-displacement products almost absent (0.03 on gap27-NCP227). PARP2 has the highest affinity for nick-NCP227 compared to those of Polβ and PARP1 (Table 1), which is fully consistent with the reaction being arrested precisely at this intermediate. PARP2-induced inhibition of the single-nucleotide gap-filling reaction was, however, detected when Polβ activity directly in this reaction was assayed in the presence of dTTP only, at a lower Polβ concentration, i.e. under conditions of a low extent of substrate conversion (below 50%; Supplementary Fig. 5). Thus, competition between PARP2 and Polβ for interaction with the 1-nt DNA gap strongly depends on the experimental conditions, and the apparent selectivity of PARP2 for the nicked intermediate reflects rather the relative preference as evidenced from data in Table 1.

**Fig. 5.**
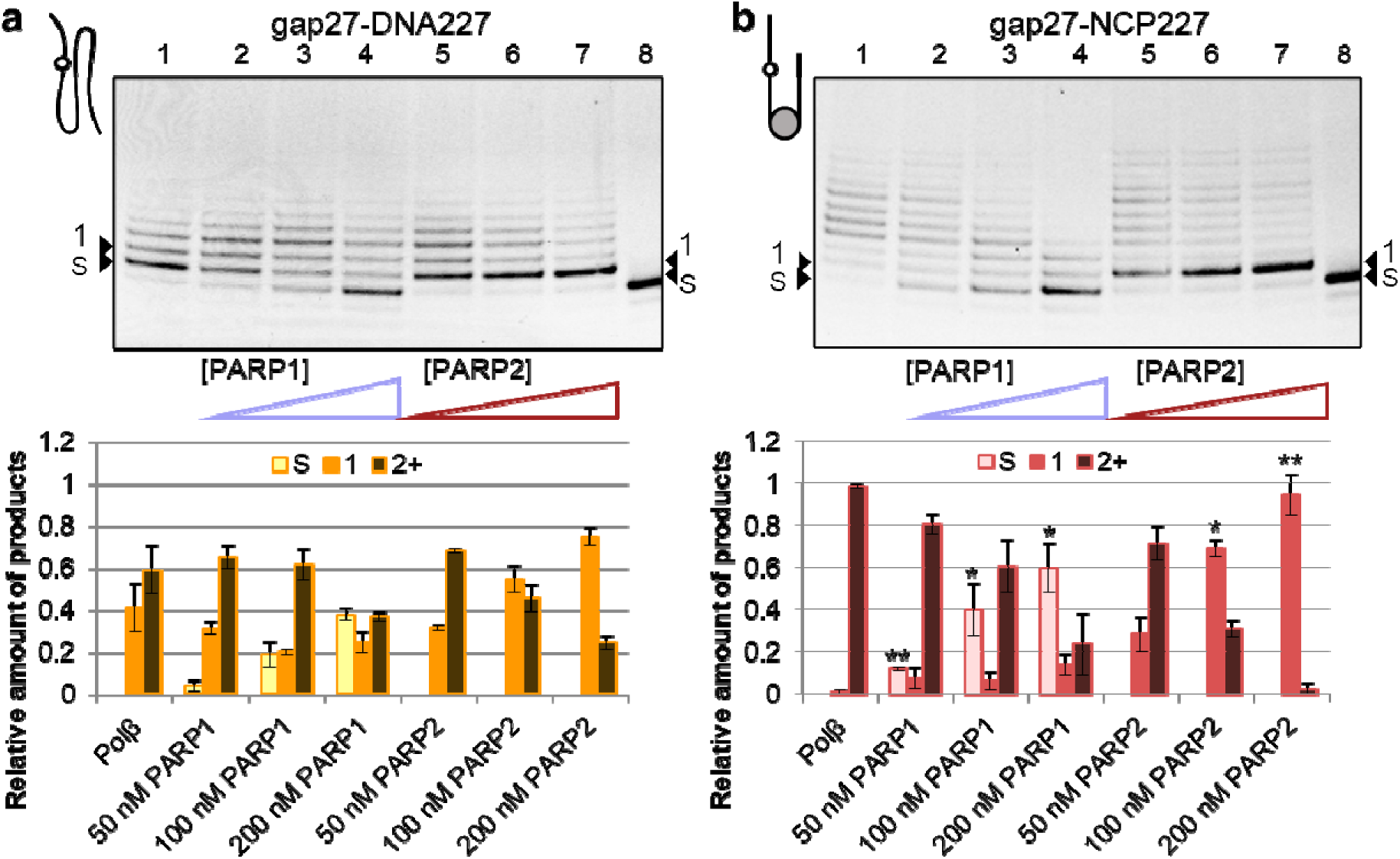
PARP1 and PARP2 inhibit distinct stages of Polβ-catalyzed DNA synthesis, acting more efficiently on the nucleosome substrate. (a, b) Electrophoregrams and quantification of DNA extension by Polβ via strand-displacement synthesis on gap27-DNA227 (**a**) or gap27-NCP227 (**b**). 50 nM substrates were incubated with 10 nM Polβ, four dNTPs (100 μM each) and increasing concentrations of PARP1 (50, 100 or 200 nM; lanes 2–4) or PARP2 (50, 100 or 200 nM; lanes 5–7) for 10 min. Lane 1 – Polβ alone; lane 8 – substrate without Polβ. Positions of the substrate (S) and of the product of single nucleotide insertion (1) in denaturing 15% PAG are indicated to the left and right of gel images. Lower panels show relative amounts of the unreacted substrate, the single-nucleotide insertion product (1) and the strand-displacement products (2+), quantified from independent experiments (mean ± SD, n = 3). Asterisks denote significant differences between the corresponding values for gap27-DNA227 and gap27-NCP227 (Student’s t-test, *p ≤ 0.05; **p ≤ 0.01).

Interestingly, at any concentration of PARP1/PARP2, the inhibitory effect on strand-displacement synthesis was more pronounced with the nucleosome substrate than with the naked DNA (Fig. 5). This difference was most evident for PARP2: at 200 nM it reduced the yield of strand-displacement products 49-fold on gap27-NCP227 versus 2.5-fold on gap27-DNA227. Lower PARP1 and PARP2 concentrations were required to suppress synthesis on gap-NCP to the same level as on gap-DNA. Although no substantial difference in the binding affinity of Polβ, PARP1, or PARP2 for gap-DNA vs gap-NCP was detected (Table 1), it is evident that the proximal nucleosome modulates the functional interplay between PARP enzymes and Polβ at the DNA lesion.

To verify that the inhibitory action of PARPs results from competition for the damage site rather than from a DNA-independent interaction with Polβ, we explored modulation of the inhibition effects by addition of an undamaged DNA competitor: linear non-gapped DNA227 or supercoiled pBR322 plasmid lacking strand breaks. The addition of non-gapped DNA227 significantly reduced the PARP1-mediated inhibition: the fraction of unreacted substrate (S) decreased approximately 1.4-fold on gap-DNA227 and 2-fold on gap-NCP227 (Supplementary Fig. 6a), whereas the PARP2-mediated inhibition remained essentially unchanged. Notably, non-gapped DNA227 carries the same DNA ends as the substrate but no internal gap or nick, so its failure to reduce the PARP2 effect indicates that this effect requires the strand break itself rather than the free DNA ends. The strength of PARP1-induced inhibition likewise decreased with increasing pBR322 concentration (Supplementary Fig. 6b), regardless of the substrate used, indicating that PARP1 was efficiently sequestered by the undamaged DNA and must be bound at the lesion to interfere with Polβ. In contrast, the inhibition by PARP2 was insensitive to the plasmid added over the same concentration range, in line with the strict requirement of PARP2 for a strand break. These observations argue that the modulation of Polβ activity by both PARPs is mediated by their binding to the BER intermediates, and additionally illustrate the distinct DNA damage preferences of the two enzymes.

### PARylation Antagonizes the Inhibitory Action of PARPs and H1 on Strand-Displacement Synthesis

PARP1 and PARP2 catalyze synthesis of poly(ADP-ribose) (PAR) attached to themselves (automodification) and target proteins, using NAD⁺ as a substrate. The autoPARylation promotes dissociation of PARPs from damaged DNA. Furthermore, these enzymes can PARylate core and linker histones in the presence of HPF1. We analyzed how addition of NAD⁺ modulates the effects of PARP1 and PARP2 on Polβ-catalyzed DNA synthesis on the nucleosome, including in the presence of H1 and HPF1 (Fig. 6, Supplementary Fig. 7a). As expected, the inhibitory action of PARP1 and PARP2 was nearly completely abolished in the presence of NAD⁺; the efficiency of strand-displacement synthesis was restored under PARylation conditions (Fig. 6a, b; compare lanes 2 and 6). Quantification of insertion products of 24-nt length and longer, i.e. those synthesized through the entire linker and further, confirmed this observation: their fraction was significantly reduced in the presence of PARP1/PARP2 and H1, alone or together, whereas under PARylation conditions it was closer to that produced by Polβ alone, both in the absence and presence of H1 (Fig. 6a, b). PARP1 suppressed the DNA synthesis most efficiently, compared to PARP2 and especially to H1, and this effect predominated when PARP1 and H1 were present together (Fig. 6a, compare lanes 3, 4 and 5). However, effects produced by PARP2 and histone H1 present together were different from those detected for each protein alone (Fig. 6b, compare lane 5 with lanes 3 and 4), indicating combined action of the two proteins. Most intriguingly, the restrictive effect on the length of extension products produced by H1 was not observed under PARylation conditions (Fig. 6a, b; compare lane 4 with lane 7). This fact was also observed in the presence of HPF1 (Supplementary Fig. 7a, b). NAD⁺ itself was without effect in the absence of PARP (Supplementary Fig. 7a, lane 6), i.e., the decrease in the inhibition effect of H1 requires PAR synthesis. Additional experiments show that the inhibitory action of PARP1/PARP2 and H1 under PARylation conditions depends on the NAD^+^ concentration known to control the efficiency of PAR elongation (Supplementary Fig. 7b). The dose-dependent nature of this effect provides insight into the mechanism of inhibition: given that autoPARylation primarily promotes the release of PARPs from DNA, the gradual recovery of synthesis with increasing NAD⁺ indicates that both PARPs must remain bound to the damaged site to interfere with Polβ activity. Taken together with the sequestration of PARP1 by DNA competitors described above (Supplementary Fig. 6), this supports the competition for the BER intermediate as the reason of the observed inhibition. These results suggest that the activity of PARP1 and PARP2 is sufficient to displace H1 from the nucleosome entry/exit site. The eviction could occur either through direct PARylation of H1 or via its non-covalent interaction with PAR^58^. To clarify the mechanism, we further analyzed the PARP1/PARP2 catalyzed modification of H1.

**Fig. 6.**
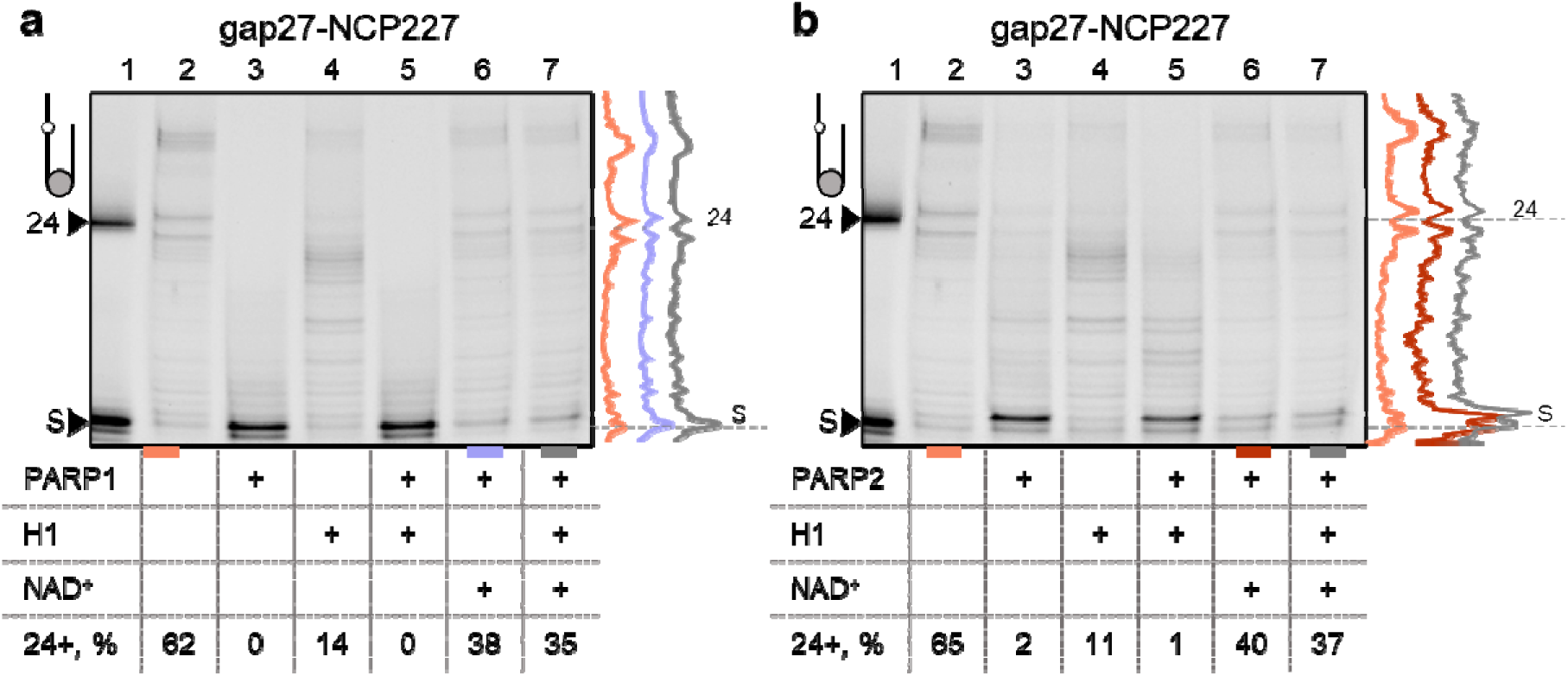
Influence of H1 and PARylation on Polβ activity. (a, b) Electrophoregrams show DNA extension by Polβ via strand-displacement synthesis after incubation of 50 nM Polβ with 50 nM gap-NCP, four dNTPs (100 μM each), without or with addition of 75 nM H1, 100 nM PARP1/PARP2, and 100 μM NAD. PARylation was carried out by preliminary incubation of gap-NCP mixtures with PARP1/PARP2 and NAD, without or with H1, for 30 min; fur her incubation with Polβ was performed for 10 min. Lane 1 – control indicating the positions of substrate and 24-mer extension product. On the right are curves reflecting relative intensities of bands in samples compared (marked with color under the electrophoregram). The percentage of products longer than 24 nucleotides (24+) is indicated below each lane.

### Linker histone H1 is modified by PARP1 and PARP2 in an HPF1-dependent manner

The PARylation-dependent effect of histone H1 on Polβ activity prompted us to explore the direct PARylation of H1; a key question was whether the covalent modification of H1 itself is sufficient to account for its release from the nucleosome entry/exit site under our experimental conditions, in which the HPF1 factor was absent. In early studies, H1 was described as an acceptor of ADP-ribose modified at glutamate residues^59,60^, whereas more recent work has shown that ADP-ribosylation of histones, including H1, is strongly stimulated by HPF1^37,61^. Thus, we verified which of these two modes of modification operates with the gap-NCP substrate and the recombinant H1.0 preparation used in this study. The PARylation reaction was performed using gap-NCP for PARP1/PARP2 activation in the presence and absence of HPF1 (Fig. 7). As established previously^16,17,45^, PARP1 was more active in the automodification than in histone modification, whereas PARP2 catalyzed ADP-ribosylation of histones more efficiently (Fig. 7a). The presence of H1 reduced the level of HPF1-dependent modification of core histones by both enzymes due to redistribution of PARylation targets between the linker and core histones (Fig. 7a, c; compare samples 4, 8 with samples 3, 7). In the absence of HPF1, modification of H1 was detected only at its increased concentration as evidenced from analysis of ADP-ribosylation products after treatment of reaction mixtures with poly(ADP-ribose) glycohydrolase (PARG) (Fig. 7b, c and Supplementary Fig. 8). Thus, the very low level of HPF1-independent PARylation of H1 by PARP1/PARP2 is most likely insufficient to induce the linker histone eviction. Additional fluorescence anisotropy experiments showed that while H1 slowed down the kinetics of PARylation-induced dissociation of PARP1/PARP2 from the complex with gap-NCP, it did not affect the final level of anisotropy (Supplementary Fig. 9), suggesting that H1 dissociates from the complex together with PARPs. Therefore, the eviction of H1 is mediated by its high-affinity interaction with PAR (demonstrated previously^58,62^) rather than by direct PARylation (which is minimal without HPF1). We propose that the automodification of PARP1 and PARP2 leads to the eviction of H1 from the nucleosome due to its attraction to PAR, ultimately enabling strand-displacement synthesis to proceed through the entry/exit region into the nucleosome.

**Fig. 7.**
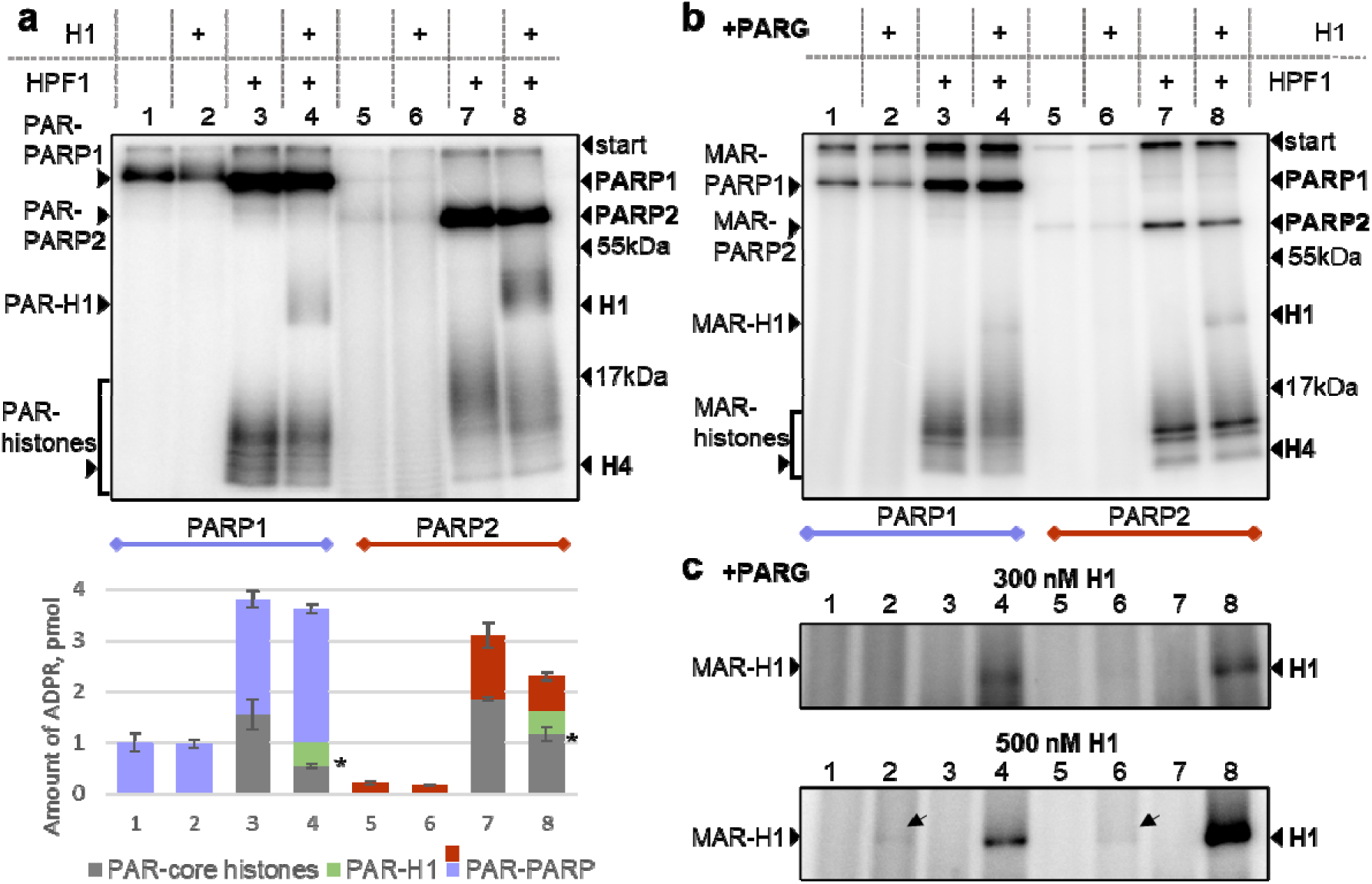
PARylation of histone H1 depends on HPF1. (a, b) Autoradiograms show covalent binding of P-labeled PAR or MAR to protein targets after incubation of PARP1/PARP2 (500 nM) with [ P]NAD (1 μM) and gap-NCP (250 nM) for 30 min, without or with addition of H1 (70 nM) and HPF1 (1 μM) before (**a**) and after (**b**) subsequent treatment of reaction mixtures with PARG. Positions of ADP-ribosylated proteins and their native forms (and molecular weight markers) in 20% SDS-PAG are indicated to the left and right of gel images. The histogram shows the amount of PAR attached to PARP1/PARP2, H1 and histones in the distinct samples (the mean ± SD, n=3). Asterisks denote significant differences in the level of core histone modification in the presence versus absence of H1 (samples 3 vs 4 and 7 vs 8; Student’s t-test, p ≤ 0.05). (**c**) - Autoradiograms show covalent binding of P-labeled MAR to H1 after incubation of 500 nM PARP1/PARP2 with 1 μM [P]NAD, 250 nM gap-NCP, 1 μM HPF1 and 300 nM or 500 nM H1 for 45 min and following PARG treatment. Positions of mono-ADP-ribosylated H1 and its native form in 20% SDS-PAG are indicated to the left and right of gel images.

Interestingly, the yields of PARylated core histones were significantly different in the absence and presence of H1, but the yields of MARylated products were practically independent of the presence of H1 (compare lanes 2, 4, 7 and 8 in Fig. 7a with respective lanes in Fig. 7b). These data suggest that H1 inhibits modification of core histones most likely at the elongation stage. This may be attributed to different stabilities of complexes formed by the linker histone and core histones with DNA, or it may point to an interaction between H1 and PAR that may influence the PAR chain length, as shown previously for other PAR readers^63,64^.

## Discussion

The canonical view of chromatin as a passive barrier to DNA repair has been increasingly refined to encompass its role as an active regulator of repair pathway choice^19,28,49^. While it is well established that the nucleosome core particle (NCP) profoundly suppresses the activity of BER enzymes, most studies have focused on lesions buried within the nucleosomal DNA and the regulatory landscape of the linker DNA has remained less explored^49^. Whereas a lesion within the core DNA is subject to severe steric hindrance^14,21–24,40^, a lesion in the linker is physically accessible yet its processing is influenced by the nearby NCP. Here, we investigate DNA repair synthesis in the linker DNA and reveal how the adjacent NCP, in conjunction with associated proteins, creates a unique regulatory environment that modulates the activity of Polβ and the balance between the short-patch (SP) and long-patch (LP) sub-pathways of BER.

Our most striking finding is that NCP can stimulate Polβ-catalyzed single-nucleotide gap-filling and strand-displacement synthesis on adjacent linker DNA, instead of expected suppression. The effect is specific to this mechanism of DNA repair synthesis: the observed rate constants of single-nucleotide gap filling differed no more than 1.6-fold between the naked and the nucleosome substrates, whereas the distribution of strand-displacement products shifted towards the long products by several fold. Two observations constrain the possible explanation. First, the stimulation is not due to a stronger binding of the enzyme, since the affinity of Polβ for gap27-NCP227 was only 1.5-fold higher than that for gap27-DNA227 – a difference comparable to the variation in the gap-filling rates but insufficient to explain the shift in product distribution. Second, and more importantly, a non-gapped NCP present as an extrinsic factor stimulated the repair synthesis on the naked gapped DNA, which excludes any mechanism requiring the particle to be positioned next to the lesion on the same DNA molecule. The inability of free (unassembled) core histones to produce such effect and absence of stable complexes of Polβ with the non-gapped NCP (not detected by either mass photometry or EMSA) argue that the nucleosome acts neither through nonspecific enzyme stabilization nor through direct binding to the polymerase. We therefore favor a model in which the nucleosome acts on the displaced DNA strand. Gap filling generates no 5’-flap, whereas strand-displacement does; a particle able to engage the single-stranded flap would shift the local equilibrium away from its re-annealing and thereby favor continued displacement synthesis. Such a model accounts for the specificity of the effect for the strand-displacement mechanism, for its proceeding by nucleosome acting less efficiently in trans than in cis (i.e. as the inherent part) since the last variant ensures a constant presence of the factor in the region of action. We emphasize that no direct interaction between the nucleosome and the flap has been demonstrated here; the model represents an interpretation consistent with the entire data obtained.

This stimulatory effect extends beyond the action of Polβ alone. Our data reveal that the presence of an adjacent nucleosome not only influences Polβ activity but also orchestrates its functional interplay with the downstream enzyme of the LP-BER sub-pathway, FEN1. The rhythmic pattern of extension products, where bands corresponding to every third nucleotide are faint, detected specifically on the nucleosome substrate, is highly indicative of effective substrate channeling, probably via the "passing the baton" mechanism^5^. Notably, this coupling, unlike the stimulation of strand-displacement synthesis itself, was not reproduced by a nucleosome acting in trans: an added NCP increased the overall length of the products on gap27-DNA227 without inducing the periodic pattern, and attenuated it on gap27-NCP227 (Supplementary Fig. 3).

Consistent with this model, the interaction of PARP1, Polβ and FEN1 at the branch-point BER intermediate has been directly demonstrated by their co-precipitation following photocrosslinking to this DNA intermediate in cell extracts^65,66^. Importantly, this coordination on the nucleosome contrasts with the more stochastic and asynchronous product distribution on naked DNA, indicating that the nucleosome context actively facilitates efficient handoff between the repair enzymes. In addition, the inhibitory effects of both PARP1 and PARP2 (discussed below) were more pronounced on the nucleosome substrate than on the naked DNA, despite similar binding affinities. It has been shown previously that the protein assembly for a specific repair process can be coordinated by protein-protein interactions as has been determined for the core enzymes of BER^66–69^. The Polβ–FEN1 coordination thus requires the nucleosome positioned next to the lesion in cis, which supports its role as a localized assembly platform rather than a diffusible stimulator. Here we propose that the nucleosome itself functions as a platform that fosters coordinated substrate channeling in the linker DNA region. The notion that the nucleosome is an active partner rather than a passive obstacle for the BER machinery is in line with structural studies that have resolved how BER enzymes engage nucleosomes^24,30,33,70–73^. Furthermore, we demonstrate that Polβ forms higher-order complexes on gapped DNA and nucleosome substrates via multimerization. The propensity for self-association has been previously noted for many BER players, including PARP1 and PARP2^17,64,68^. Binding of several Polβ molecules to gapped and template-primer structures was shown^74^. This observation raises the possibility that multiple cycles of Polβ binding and flap endonuclease-catalyzed cleavage could be coordinated through such multimeric assemblies.

The stimulatory effect of the nucleosome is balanced by the suppressive action of the linker histone H1, which in turn is controlled by PARP-catalyzed PAR synthesis. The linker histone H1 contributing to chromatin compaction^52–54^ limits Polβ-catalyzed DNA repair synthesis at the entry/exit site of the nucleosome. It is noteworthy that this restriction is spatially localized: products accumulate as the synthesis approaches the position corresponding to the boundary between the linker and the core – precisely where H1 is located. Critically, this restriction is reversed by PARP1/PARP2-catalyzed PARylation. While eviction of histone H1 *in vivo* and its function as a high-affinity PAR-reader *in vitro* have been established^51,58,62,75,76^, the dominant mechanism of H1 by which PARylation evicts H1^77^. Our data show that H1 eviction from DNA occurs even when its direct PARylation (detected in the absence of HPF1) is minimal, favoring a model where H1 is sequestered via PAR-mediated interaction with the automodified PARP (Fig. 8). The same conclusion is supported by an independent approach: H1 slowed the release of PARP1 and PARP2 from the nucleosome substrate without affecting the final level of dissociation curves (Supplementary Fig. 9); this is consistent with the expected scenario in which the linker histone leaves the complex together with the automodified enzyme rather than being released independently. Taken together, these observations point to a dynamic, PARylation-controlled mechanism for primary chromatin relaxation required for DNA repair.

**Fig. 8.**
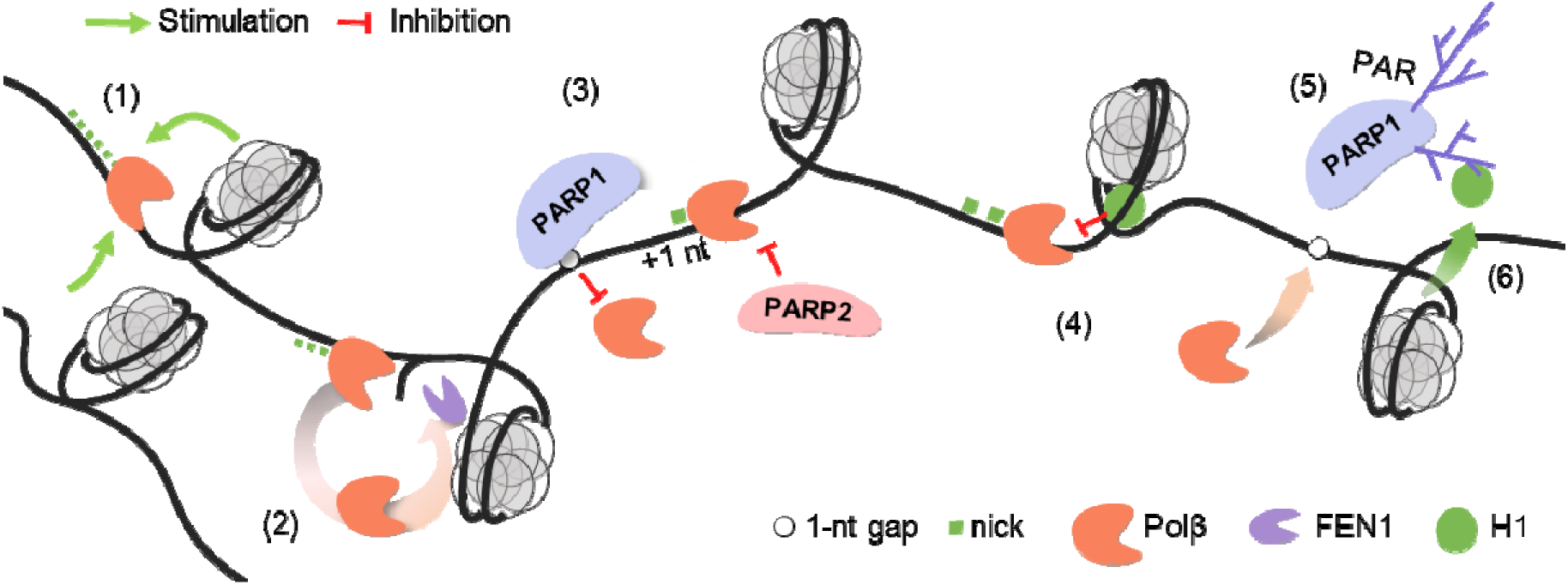
A model of multi-layered regulation of Polβ activity in a nucleosome context. (1) The adjacent nucleosome stimulates Polβ strand-displacement synthesis in the linker DNA; (2) The nucleosome coordinates the handoff between Polβ and FEN1, promoting coordinated synthesis/cleavage cycles (substrate channeling); (3) PARP1 and PARP2 suppress repair synthesis at different steps: PARP1 competes for the 1-nt gap, preventing the first insertion, whereas PARP2 acts at the resulting nick, arresting strand-displacement after the first nucleotide (+1 nt); (4) Linker histone H1 bound at the nucleosome entry/exit site restricts strand-displacement as synthesis approaches the boundary; (5) PARP1/PARP2-catalyzed autoPARylation releases the PARPs from DNA, reducing their inhibition of Polβ; (6) the autoPARylation of PARPs also evicts histone H1 from the entry/exit region, thereby modulating its effect.

Our findings refine the distinct roles of PARP1 and PARP2 in BER. Both PARPs are well established to suppress strand-displacement synthesis via competition with BER enzymes for DNA intermediates^12,13,40,67,78^, and their dissimilar preferences for the distinct BER intermediates, including the higher affinity of PARP2 for nicked DNA, have been documented by biochemical, single-molecule and, more recently, biophysical approaches^13,14,17,79^. These preferences coupled with the stage-specific inhibitory effects of PARPs, suggest a functional specialization: PARP1 appears to modulate predominantly the initial stages of BER, while PARP2 exerts a major impact on the following steps (DNA synthesis and ligation). In our experiments, both enzymes compete with Polβ for substrate binding, but their modes of inhibition are distinct. PARP1 suppresses 1-nt gap-filling and strand-displacement synthesis, consistent with its high affinity for the initial gapped intermediate: unreacted substrate accumulated with increasing PARP1 concentration, indicating that the very first insertion event is prevented. In contrast, PARP2, with its pronounced preference for the nicked DNA, specifically and potently inhibits the strand-displacement and is less effective at the stage of the first nucleotide insertion: no unreacted substrate was detected at any PARP2 concentration, while the single-nucleotide insertion product accumulated almost quantitatively, i.e. the reaction was arrested precisely at the nick. Two observations confirm that these effects arise from competition for the BER intermediate rather than from an interaction with Polβ in solution: the inhibition was reduced by NAD⁺ in a dose-dependent manner (since automodification releases the enzymes from the enzyme-DNA complex), and the inhibitory effect of PARP1 (though not PARP2) was abolished by the addition of DNA competitor. Thus, the DNA synthesis step itself defines the functional boundary between the regulatory domains of PARP1 and PARP2 within the BER sub-pathway. This assigns PARP2 the role of a key factor determining the choice between the SP-BER and LP-BER sub-pathways. By "blocking" the single-strand break (nick) following gap filling, PARP2 prevents strand-displacement, thereby shifting the repair process toward the short-patch (SP-BER) sub-pathway; this function is critical for linker DNA, which lacks the physical constraints inherent to the nucleosome core.

Importantly, the ability of PARP2 to bind and protect DNA intermediates containing single-strand breaks provides a mechanistic basis for the protein’s observed involvement in mediating the cytotoxic effects of PARP inhibitors. Anemia is among the most common adverse effects of PARP inhibitor therapy, and is thought to arise from the retention of PARP on unligated Okazaki fragments in rapidly dividing erythroblasts, which provokes replicative stress and cell death^80^. Genetic evidence indicates that the deleterious event is the retention of the enzyme rather than the loss of PAR synthesis: mice expressing the catalytically inactive PARP1 E988A or PARP2 E534A variants die during embryogenesis, whereas the corresponding knockouts are viable^81,82^. The two enzymes are nevertheless not equivalent in this respect, since the phenotype of the inactive PARP2 is erythroid-specific – it provokes replicative stress in rapidly dividing erythroblasts through binding to single-strand breaks on the lagging strand, impairing erythropoiesis and causing Tp53-dependent anemia^82–84^ – which matches the clinical picture and has motivated the development of PARP1-selective inhibitors with reduced hematological toxicity^85,86^. Our data provide a biochemical counterpart to this assignment: among the proteins examined here, PARP2 shows the highest affinity for the nicked intermediate and arrests DNA synthesis precisely at this step, whereas PARP1 acts one step earlier, at the gap. It should be noted, however, that selectivity of inhibition does not necessarily translate into selectivity of retention, as PARP2 can be trapped even by the PARP1-selective compound AG14361^87,88^. This distinction reinforces the need to characterize the retention of each enzyme on defined repair intermediates directly, rather than to rely on inhibition constants alone.

Thus, our study elucidates a multi-layered system to regulate Polβ activity at the crucial junction between the nucleosome core and the linker DNA. The different affinities of PARP1 and PARP2 for distinct BER intermediates create a regulatory cascade that determines the choice between short- and long-patch repair. The nucleosome can act as an active participant that influences the organization of repair factors, their kinetics, and the processing of DNA damage.

## Materials and methods

Recombinant wild-type human PARP1 and murine PARP2 were purified as described previously^89^. Recombinant wild-type human HPF1 and H1.0 were purified as described previously^45^. Human APE1, FEN1 and rat Polβ were purified as described previously^90–93^. All proteins were tested for DNA polymerase activity (Supplementary Fig. 10a). *E. coli* uracil-DNA glycosylase (UDG) and *T. aquaticus* DNA polymerase were from Biosan (Novosibirsk, Russia). Core histones were isolated from *Gallus gallus* erythrocytes and purified as described previously^94^. pGEM-3z/603 was a gift from J. Widom (Addgene plasmid #26658; http://n2t.net/addgene:26658; RRID: Addgene_26658). Synthetic oligodeoxyribonucleotides were purchased from Genterra (Moscow, Russia). The ^32^P-labeled NAD^+^ was synthesized enzymatically as described previously^95^, using [α-^32^P]ATP (with specific activity of 3000YCi/mmol, synthesized in the Laboratory of Biotechnology, ICBFM, Novosibirsk, Russia). NAD^+^ and reagents for electrophoresis and basic components of buffers were purchased from Sigma–Aldrich (USA). Double-distilled water was used to prepare all buffer solutions and reaction samples. Sequences of the oligonucleotides are presented in Table 2.

**Table 2.**
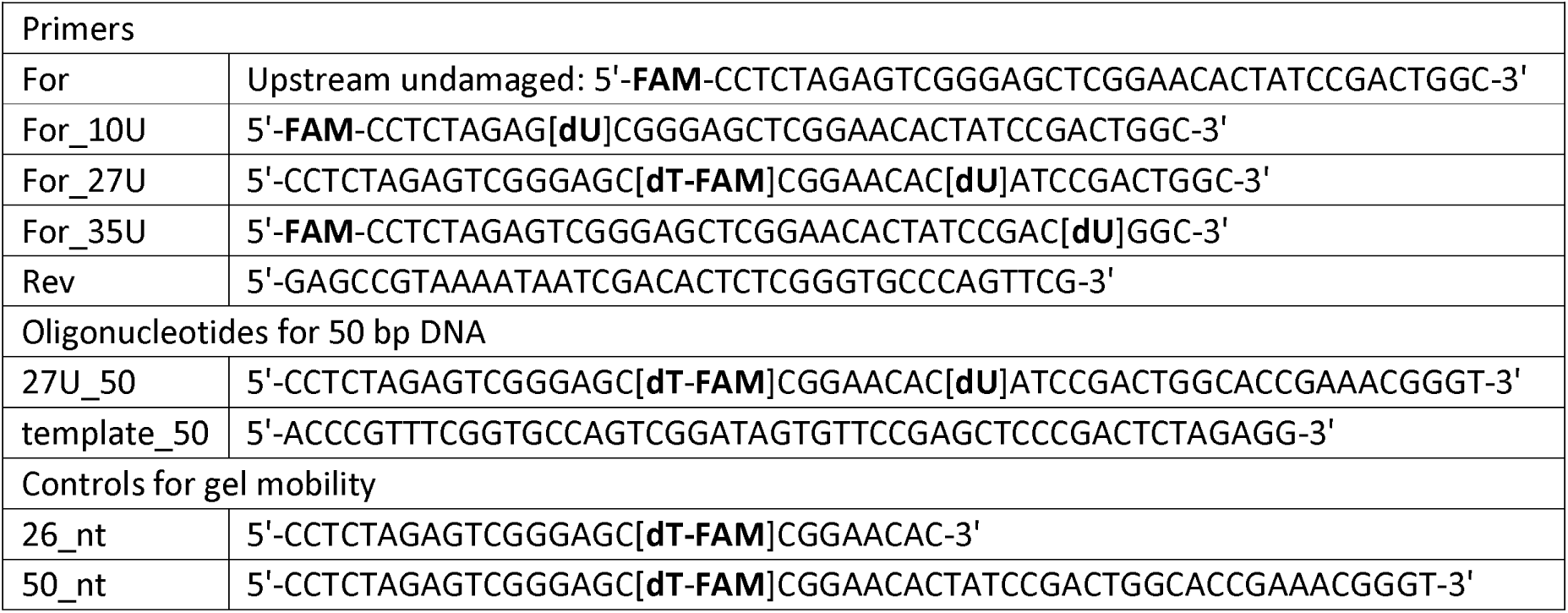
Sequences of oligonucleotides used in the study.

## Methods

### Preparation of DNA and nucleosomes

Model 227 bp DNAs containing the 147-bp core of the 603 nucleosome positioning sequence^96^ and linkers bearing dU at the predetermined position were generated by PCR from the pGEM-3z/603 plasmid using the respective oligonucleotide primers (Table 2). A 50-bp duplex (gap27-DNA50) equivalent in both the sequence and the gap position to the linker arm of gap27-DNA227 was prepared by annealing primer 27U_50 to the complementary strand template_50. Nucleosomes were assembled by mixing DNA and core histones in a high-salted buffer containing 2 M NaCl, dialyzed against the NaCl concentration gradient from 2 M to 65 mM during 6 h at 4 °C and then against the buffer with 10 mM NaCl overnight at 4 °C with gentle stirring. The optimal ratios of DNA and histones were determined in preliminary quick-time experiment as described in the published protocol^94^.

To generate a single-nucleotide gap in DNA substrates, reaction mixtures containing 1 μM DNA or NCP, uracil-DNA glycosylase (UDG, 1 activity unit per 0.6 pmol of DNA/NCP), and 0.1 µM AP endonuclease 1 (APE1) in reaction buffer (50 mM Tris-HCl (pH 8.0), 50 mM NaCl, 5 mM MgCl₂, 1 mM DTT) were incubated for 30 min at 37 °C. The final DNA/NCP preparations contained only trace amounts of 5’-dRP removed due to the lyase activity of APE1^97^ and spontaneous hydrolysis^98^ as shown previously^17^. To prepare nicked NCP (nick-NCP) substrate, reaction mixtures containing 200 nM gap-NCP, 4 nM Polβ and 10 μM dTTP in the reaction buffer were incubated for 10 min at 37 °C. Then the mixture was diluted with buffer (50 mM Tris-HCl (pH 8.0), 50 mM NaCl, 10 mM EDTA, 1 mM DTT) to 6 nM concentration of NCP and incubated for 10 min at 45 °C. The homogeneity of DNA/NCP preparations was analyzed by electrophoresis on a 4% nondenaturing PAG (Supplementary Fig. 10b).

### Fluorescence studies of Polβ/H1/PARP1/PARP2 interaction with DNAs/NCPs

The affinity of proteins for DNA and NCP substrates was evaluated by determining the half-maximal effective concentration (EC₅₀) of the protein bound to the FAM-labeled DNA or NCP using fluorescence anisotropy measurements^99^. Briefly, reaction mixtures containing 3 nM of gap-DNA (227 bp), gap-NCP, or nick-NCP and PARP1/PARP2/H1/Polβ at varied concentrations (0–400 nM) in binding buffer (50 mM Tris-HCl pH 8.0, 50 mM NaCl, 10 mM EDTA, 1 mM DTT) were prepared on ice in a 384-well plate and incubated for 5 min at room temperature. Fluorescence anisotropy measurements were performed using a CLARIOstar microplate reader (BMG Labtech). Fluorescent probes were excited at 495 nm, and emission was detected at 520 nm. Each measurement consisted of 50 flashes per well, and fluorescence intensity values were averaged automatically. Measurements for each well were performed three times at 1-min intervals. The averaged values were used for final plotting, and EC₅₀ values were calculated using the SMART Control Data Analysis software (BMG LABTECH). The data were plotted (F vs C) and fitted to four-parameter logistic equation: F = F + (F - F)/[1 + (EC_50_/C)^n^], where F is the measured fluorescence anisotropy of a solution containing the labeled DNA at a given concentration (C) of proteins, F0 is the fluorescence anisotropy of solution of the labeled DNA alone, F∞ is the fluorescence anisotropy of the labeled DNA saturated with the protein, EC50 is the protein concentration at which F - F0 = (F∞ - F0)/2, and n is the Hill coefficient, which denotes the steepness (slope) of the nonlinear curve. The experiments were performed in independent duplicates; the EC50 values are presented in Table 1.

### Mass Photometry measurements of Polβ interaction with gap-NCP substrate

The stoichiometry of Polβ complexes with gap27-NCP227 was assessed by mass photometry using a TwoMP mass photometer (Refeyn, UK). Briefly, 10 μl buffer (50 mM Tris-HCl (pH 8.0), 50 mM NaCl, 5 mM MgCl2, 1 mM DTT) was loaded to the sample chamber and the objective was focused by autofocus function. Then 0.5 μl sample containing 60 nM gap27-NCP227 (or non-gapped NCP for comparison) alone or together with 180 nM or 300 nM Polβ was added to the chamber and mixed by pipetting. The buffer was preliminary ultrafiltered by using Sartorius vivaspin columns with 1000 kDa pore PES membrane filter. The reaction mixture was prepared on ice and incubated for 5 min at room temperature. Data were collected for 1 min using the AcquireMP software (Refeyn, UK). All samples were measured using the expanded detection area. The mass photometry signals were calibrated using BSA (69 kDa) (Sigma-Aldrich, USA), recombinant PARP1 protein (113 kDa) and recombinant Cas9 protein (158 kDa). Mass photometry data were analyzed using the DiscoverMP software (Refeyn, UK) to calculate relative molecular populations from the areas of the Gaussian peaks (representing free NCP and its complexes with one or several Polβ molecules).

### Testing of Polβ-catalyzed DNA synthesis

DNA synthesis catalyzed by Polβ was performed in 5−10 µL reaction mixtures containing 50 nM DNA/NCP substrate (specified in each experiment), 20 nM Polβ (unless stated otherwise), and either 100 µM of each of four dNTPs (for strand-displacement synthesis) or 500 nM dTTP (for gap-filling synthesis) in reaction buffer (50 mM Tris-HCl pH 8.0, 50 mM NaCl, 5 mM MgCl₂, 1 mM DTT). To explore the influence of non-gapped NCP227 on the Polβ DNA polymerase activity, mixtures containing 50 nM gap27-DNA227 or gap27-NCP227 were supplemented with 50 nM NCP227. The influence of FEN1, H1, PARP1 or PARP2 on the Polβ activity was explored by the addition of 25–75 nM FEN1, 200−300 nM H1, 50–200 nM PARP1 or 50–200 nM PARP2 respectively. The concentration of Polβ was varied from 4 nM to 100 nM in distinct experiments to optimize the extent of substrate conversion and the length of extension products. To analyze the influence of H1 and PARylation on the Polβ activity, the reaction mixtures supplemented with 100 nM PARP1/PARP2, 75 nM H1, and 100 μM NAD^+^ were preincubated at 37 °C for 30 min, before the addition of Polβ and dNTPs. The DNA-synthesis reaction was performed by incubation at 37 °C for 10 min. All reactions were terminated by adding 10 µL of 8 M urea with 20 mM EDTA, followed by heating at 80 °C for 10 min and 97 °C for 1 min. Reaction products were separated by electrophoresis on 15% denaturing polyacrylamide gels. FAM-labeled oligonucleotides of 26 nt and 50 nt length (Table 2) were used as gel-mobility controls corresponding to the length of substrate and linker. Gels were visualized using a Typhoon FLA 9500 scanner (GE Healthcare Life Sciences, USA). The relative intensities of bands corresponding to the substrate and products were quantified using Quantity One 4.6.6 software (Bio-Rad, USA) as follows: the total (raw) signal of the band of substrate or product(s) (as indicated in electrophoregrams) was quantified and the same-size background signal of gel in the respective lane was subtracted from the raw signal. All the experiments were performed in three independent replicates. Statistically significant differences were evaluated by Student’s t-test (*, *p* ≤ 0.05; **, *p* ≤ 0.01).

### Kinetic measurements of Polβ activity

The reaction of DNA synthesis via strand-displacement catalyzed by Polβ was carried out in a 25 μl reaction mixture containing 50 mM Tris-HCl, pH 8.0, 50 mM NaCl, 5 mM MgCl2, 50 nM gap27-DNA50, or gap27-DNA227, or gap27-NCP227, 20 nM Polβ and 100 µM each of four dNTPs. Reactions were initiated by addition of dNTPs and stopped after 30 s, 2 min, 5 min and 10 min of incubation at 37°C by adding 5 µL aliquot of the reaction mixture to 5 µL of 8 M urea with 20 mM EDTA. Reaction products were separated and analyzed as described in subsection above. The experiments were performed in independent triplicates.

### Protein PARylation reaction

Reaction mixtures containing 250 nM gap27-NCP227, 500 nM PARP1 or PARP2, 1 μM HPF1, H1 (at varied concentrations in the range of 70−500 nM, as specified in Figure 7 legend), and 1 µM [³²P]NAD⁺ in buffer (50 mM Tris-HCl pH 8.0, 50 mM NaCl, 5 mM MgCl₂, 1 mM DTT) were incubated for 30/45 min at 37 °C. Reactions were stopped by adding 1.2 µL of inhibitor solution (50 µM olaparib, 100 mM EDTA, 25 mM Tris-HCl pH 8.0, 25 mM NaCl, 500 µM DTT). When indicated, 1 µM PARG was added to the mixtures, followed by incubation for 1 h at 37 °C to digest PAR. Subsequently, 1.5 µL of SDS-PAG sample buffer was added, and samples were heated at 97 °C for 5 min. Reaction products were separated by 20% SDS-PAG, visualized by phosphorimaging, and quantified using a Typhoon imaging system (GE Healthcare Life Sciences) and Quantity One Basic software (Bio-Rad). The experiments were performed in independent triplicates.

## Acknowledgements

We would like to thank the entire laboratory of bioorganic chemistry of enzymes for feedback. We acknowledge Svetlana N. Khodyreva for preparation of recombinant FEN1, Anton V. Endutkin and Dmitry O. Zharkov for supporting Mass Photometry experiments. We thank the reviewers for their careful reading of the manuscript and for their constructive criticism, which prompted additional experiments and helped us substantially improve the validity and clarity of the work. The reported study was funded by the Russian state-funded project for ICBFM SB RAS № 125012300658-9 (use of shared equipment for experimental work, MP experiments) and by the Russian Science Foundation № 25-74-10025 (preparation of proteins and nucleosomes) and 25-74-30006 (biochemical experiments).

## Author information

### Affiliations

Knorre Institute of Chemical Biology and Fundamental Medicine, SB RAS, Novosibirsk, Russia

Shtanov D.M., Kurgina, T. A., Kutuzov, M. M., Naumenko, K. N., Ukraintsev A. A., Moor, N. A., Lavrik, O. I.

## Contributions

D.M.S. performed the experiments, T.A.K. performed the experiments and created the figures. M.M.K., K.N.N., A.A.U. and N.A.M. contributed to the study with protein purification and nucleosome assembling. T.A.K., N.A.M. and O.I.L. analyzed the data and wrote the manuscript. All authors reviewed the results and approved the final version of the manuscript.

## Corresponding author

Correspondence to Olga I. Lavrik

## Additional information

### Supplementary information

The online version contains supplementary material available at

