## Supplementary material for "More than just a passive brick in the wall: the nucleosome facilitates DNA polymerase β activity in linker DNA and its PARP-dependent regulation in the BER sub-pathway choice"

This PDF file includes:
**Supplementary Figures 1 to 10**
**Supplementary Notes**
**Supplementary Note 1 (related to Supplementary Figure 1): Kinetic measurements of Polβ activity.**

**Supplementary Note 2 (related to Supplementary Figure 4): Electrophoretic mobility shift assay.**

**Supplementary Note 3 (related to Supplementary Figure 8): Fluorescence studies of PARP1/PARP2 dissociation from complexes with DNA during PARylation.**

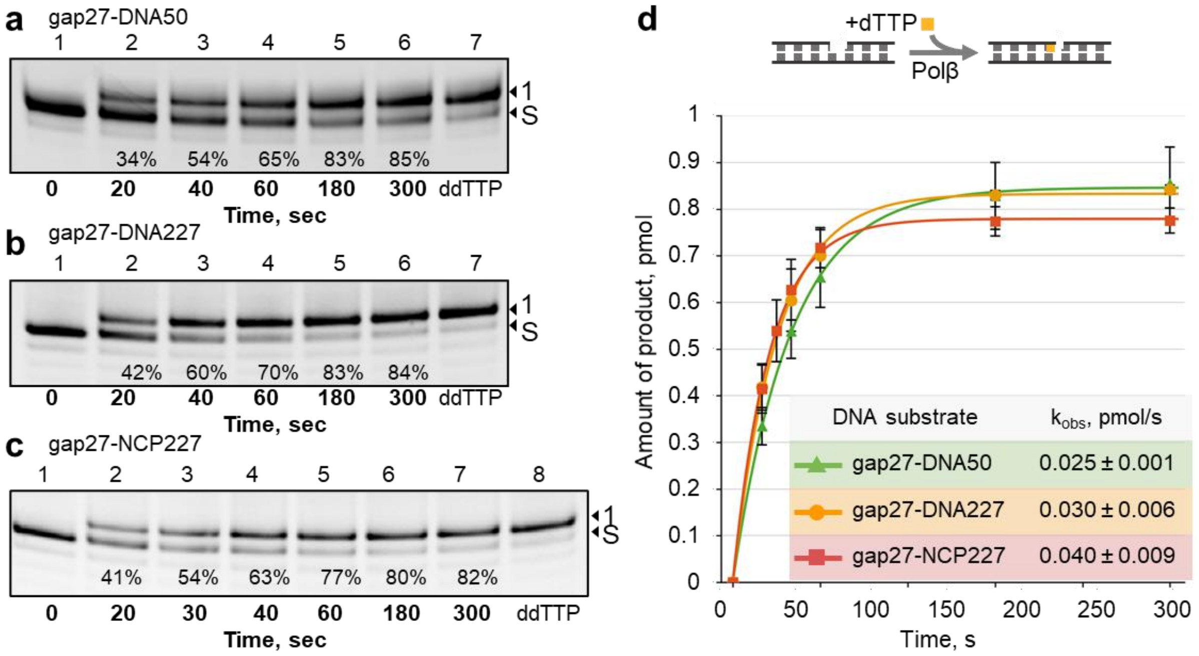


**Supplementary Fig. 1.** The rate of single-nucleotide gap-filling synthesis is similar for the naked DNA and nucleosome substrates.

(**a–c**) Time course of the gap-filling reaction catalyzed by Polβ (4 nM) on gap27-DNA50 (a), gap27-DNA227 (b) or gap27-NCP227 (c) (200 nM) in the presence of dTTP (500 nM). Reactions were stopped after the time intervals indicated below each lane; the extent of substrate conversion is given underneath. The last lane of each gel (ddTTP) shows the reaction performed with ddTTP instead of dTTP, confirming that a single nucleotide is incorporated under these conditions. Positions of the substrate (S) and of the single-nucleotide insertion product (1) in denaturing 20% PAG are indicated to the right of electrophoregrams.

(**d**) Accumulation of the gap-filling product with time for the three substrates (mean ± SD, n = 3); curves are fits to a single-exponential function. The inset lists the observed rate constants (k_obs) calculated from the fits. The differences between $k_{obs}$ values for gap27-DNA50 and gap27-DNA227 as well as for gap27-DNA227 and gap27-NCP227were statistically insignificant.

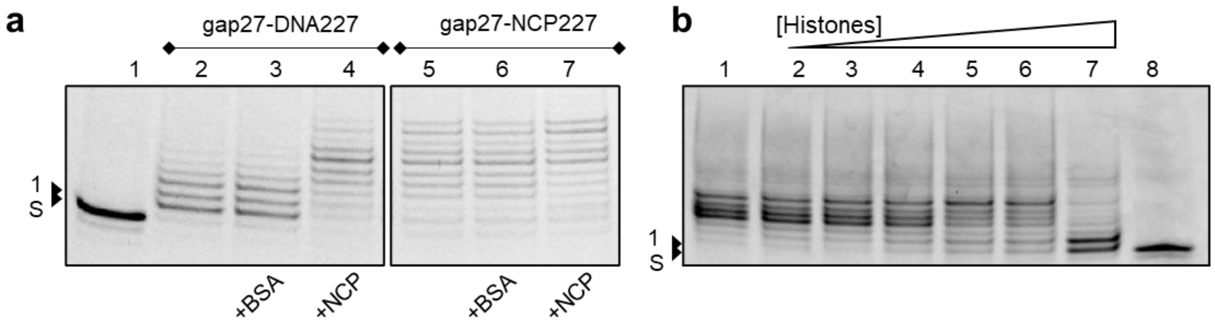


**Supplementary Fig. 2.** The stimulatory effect of the adjacent nucleosome on DNA repair synthesis in linker region is not reproduced by free core histones.

(**a**) gap27-DNA227 (lanes 1–4) or gap27-NCP227 (lanes 5–7) (50 nM) was incubated with four dNTPs (100 μM each) for 10 min. Lane 1 – substrate alone; lanes 2, 5 – with Polβ (20 nM); lanes 3, 6 – with Polβ (20 nM) and BSA (50 nM); lanes 4, 7 – with Polβ (20 nM) and non-gapped NCP227 (50 nM).
(**b**) gap27-DNA227 (50 nM) was incubated with Polβ (20 nM), four dNTPs (100 μM each) and increasing concentrations (6,25–100 nM) of free core histones (lanes 2–7) for 10 min; lane 1 – without histones, lane 8 – without Polβ.
Positions of the substrate (S) and of the single-nucleotide insertion product (1) in denaturing 15% PAG are indicated to the left of the gel images.

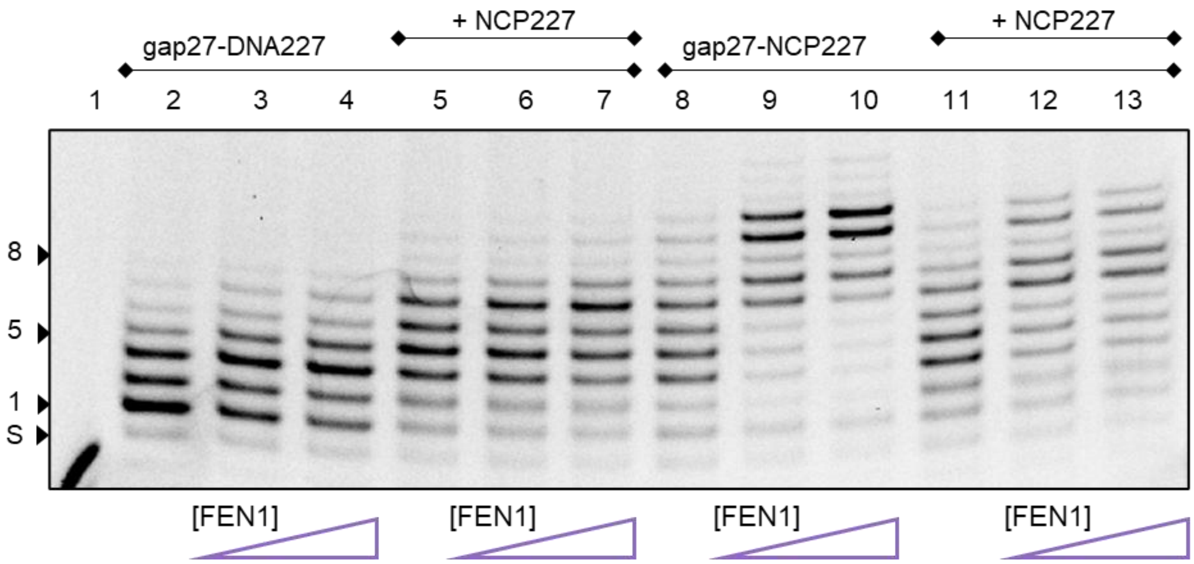


**Supplementary Fig. 3**. Influence of the intact nucleosome on interaction of Polβ and FEN1.

gap27-DNA227 (lanes 2–7) or gap27-NCP227 (lanes 8–13) (50 nM) was incubated with 16 nM Polβ and four dNTPs (100 µM each) for 10 min without FEN1 (lanes 2, 5, 8, 11), with 12,5 nM FEN1 (lanes 3, 6, 9, 12) or with 25 nM FEN1 (lanes 4, 7,10, 13) in the absence (lanes 2–4 and 8–10) or presence (lanes 5–7 and 11–13) of 50 nM non-gapped NCP227. Positions of the substrate (S) and of products with 1, 5 and 8 inserted nucleotides (1, 5 and 8) in denaturing 15% PAG are indicated to the left of the gel image.

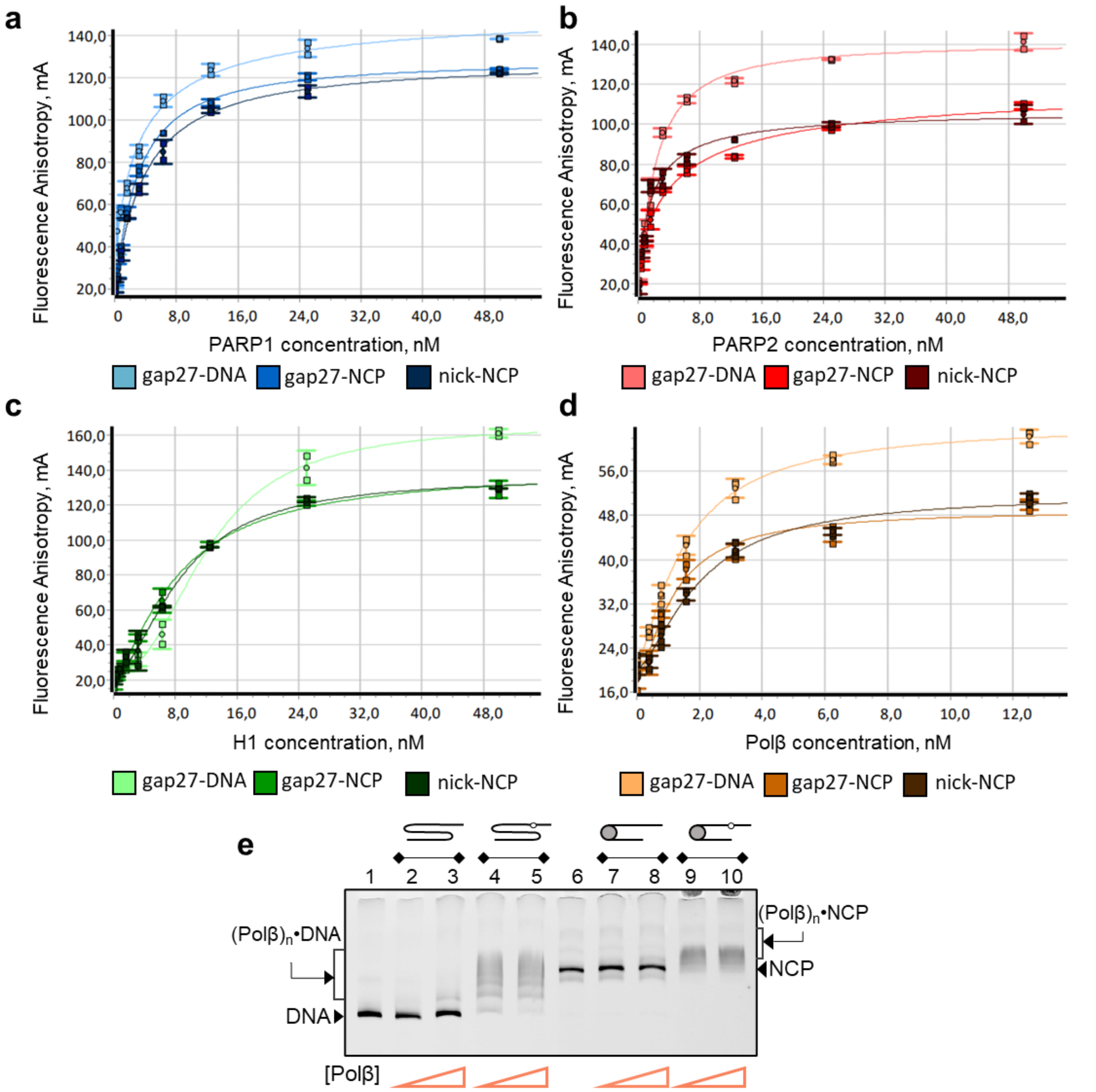


**Supplementary Fig. 4.** Binding of Polβ, PARP1, PARP2 and H1 to the gapped and nicked DNA and nucleosome substrates.

(**a–d**) Typical titration curves detected by measuring fluorescence anisotropy of FAM-labeled gap27-DNA227, gap27-NCP227 and nick-NCP227 (3 nM) in the presence of increasing concentrations of PARP1 (**a**), PARP2 (**b**), H1 (**c**) or Polβ (**d**). Data points represent the mean ± SD of three parallel measurements; curves are fits to the four-parameter logistic equation used to derive the $EC_{50}$ values listed in Table 1.

(**e**) Complex detection by EMSA. Electrophoregram of 5% native PAG shows complex formation by increasing concentrations (50–200) of Polβ with 50 nM undamaged DNA227 (lanes 2, 3), 50 nM gap27-DNA227 (lanes 4, 5), 50 nM undamaged NCP227 (lanes 7, 8) and 50 nM gap27-NCP227 (lanes 9, 10); lanes 1 and 6 – DNA227 and NCP227 alone. Higher-order complexes, $(Pol\beta)_n$•DNA/$(Pol\beta)_n$•NCP, are formed with the gapped substrates only. Positions of free DNA, free NCP and the complexes are indicated.

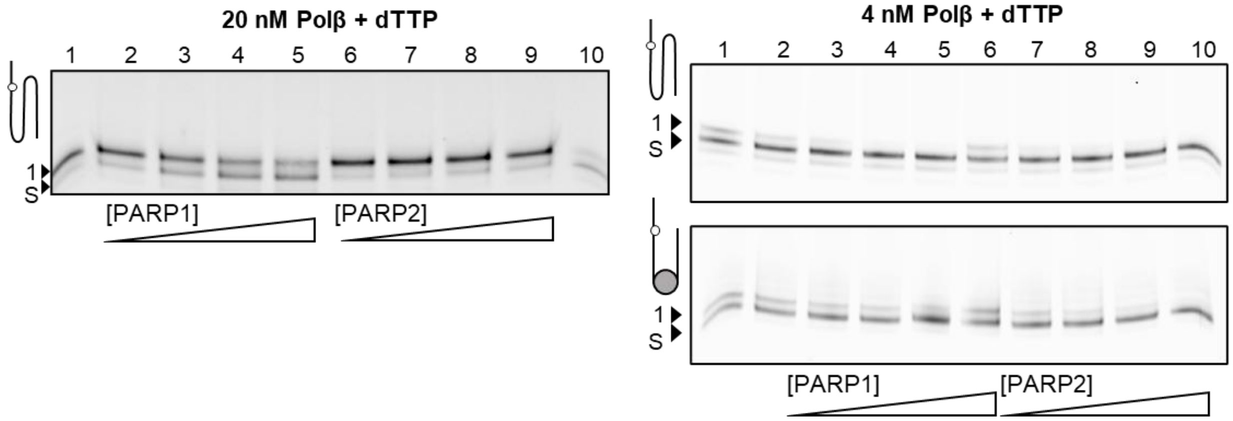


**Supplementary Fig. 5.** PARP1- and PARP2-induced inhibition of gap-filling reaction depends on the concentration of Polβ.

The single-nucleotide gap-filling reaction was performed using gap-27-DNA227 or gap-27-NCP227 as a substrate (indicated to the left of each gel image) by incubation with 20 nM or 4 nM Polβ and dTTP, in the absence (lane1) and presence of increasing PARP1/PARP2 concentration (100–400 nM) for 3 min. At the lower Polβ concentration, where substrate conversion remained below 50%, inhibition by PARP2 became detectable.

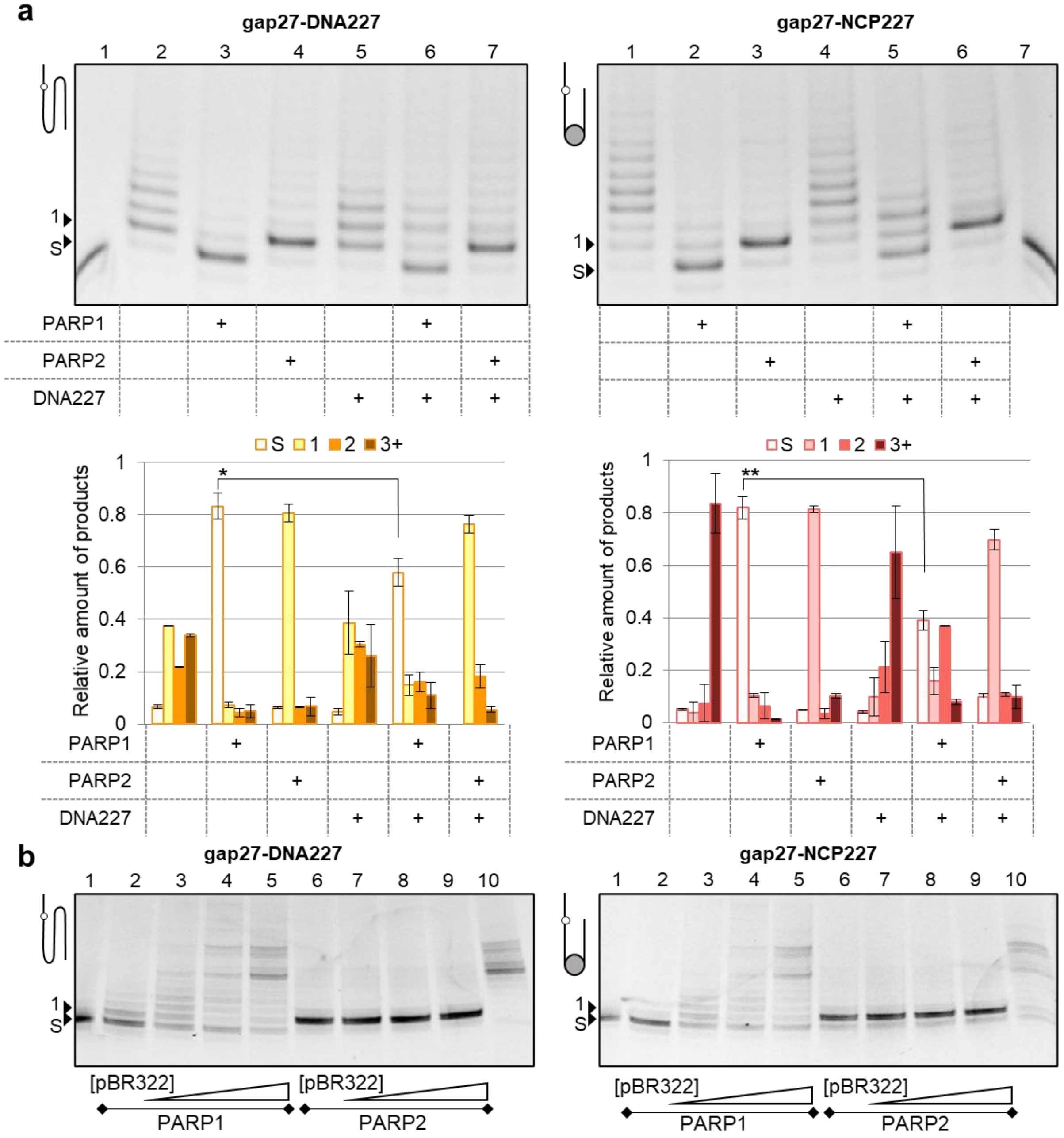


**Supplementary Fig. 6. Impact of an intact DNA on PARP-induced inhibition of Polβ activity.**

(**a**) The inhibition effect of PARP1/PARP2 decreased in the presence of non-gapped DNA227. Each of two substrates, gap27-DNA227 or gap27-NCP227, (50 nM) was incubated with 20 nM Polβ (lanes 2-7, left image; lanes 1-6, right image) and dNTPs (100 µM each) in the absence or presence (+) of 100 nM PARP1/PARP2), without or with addition of 50 nM DNA227. Line 1 for gap27-DNA227 and lane 7 for gap27-NCP227 – substrate without Polβ.

(**b**) Supercoiled pBR322 plasmid affects the inhibitory action of PARP1. Each of two substrates, gap27-DNA227 or gap27-NCP227, (50 nM) was incubated with 20 nM Polβ and dNTPs (100 µM each) in the presence of 200 nM PARP1/PARP2 (lanes 2–5/6–9), without (lanes 2 and 6) or with addition of increasing concentrations (5–20 nM) of pBR322. Lane 10 – incubation with Polβ without PARP1/PARP2 and pBR322. The inhibition by PARP1 decreased in a dose-dependent manner, whereas that by PARP2 is independent of the pBR322 addition.

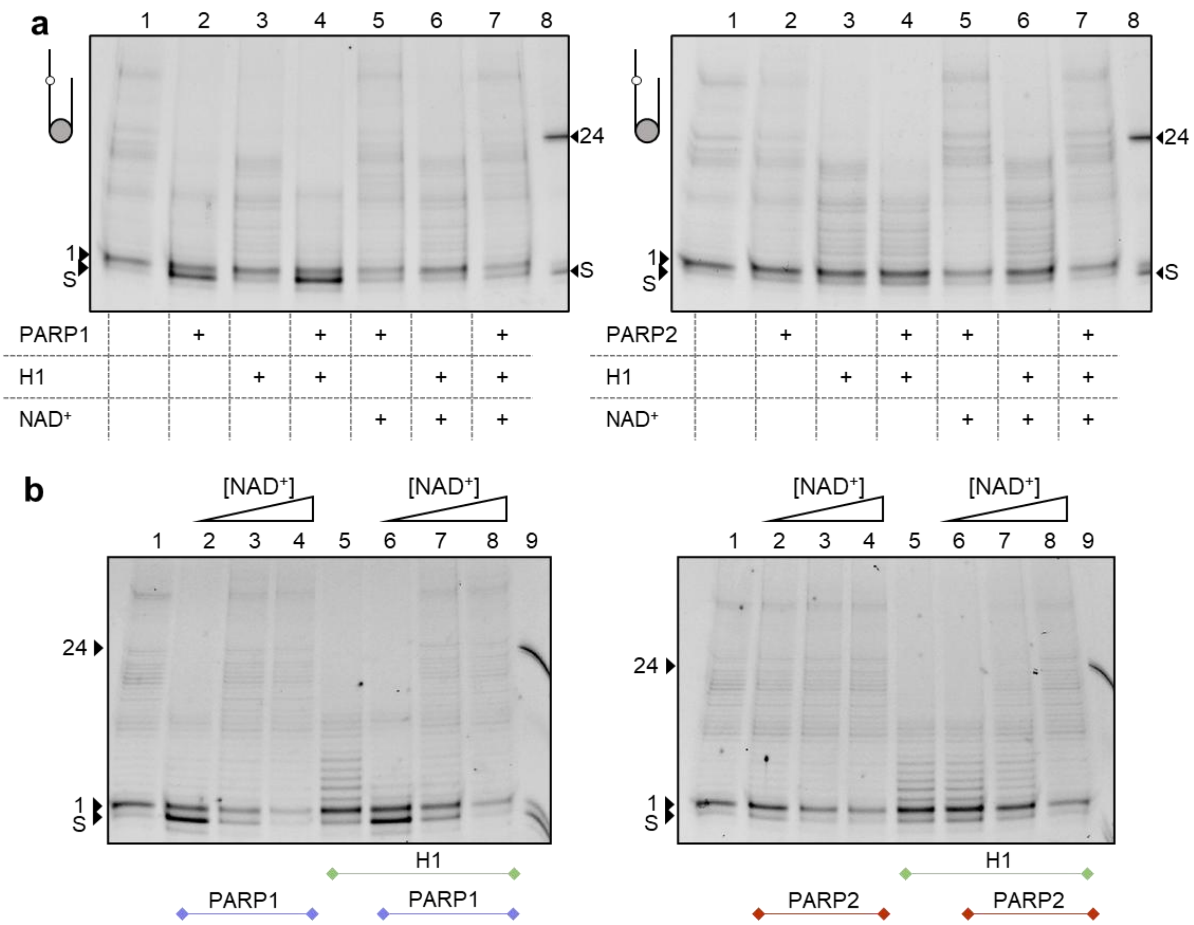


**Supplementary Fig. 7.** Effects of HPF1 and $NAD^+$ concentration on the PARP- and H1-mediated inhibition of Polβ activity.

(**a**) Strand-displacement synthesis by Polβ (50 nM) on gap27-NCP227 (50 nM) in the absence or presence (+) of 100 nM PARP1/PARP2, 75 nM H1 and 100 µM $NAD^+$, as indicated below each gel, with HPF1 present in all samples. Lane 8 – control lane to show the positions of the substrate and 24-mer product in denaturing 20% PAG. The H1-mediated restriction of the product length was not observed under conditions of HPF1-dependent PARylation.

(**b**) The same reaction was carried out in the absence (lanes 1, 5) or presence of PARP1/PARP2 (lanes 2-4, 6-8), without (lanes 1-4) or with addition of 75 nM H1 (lanes 5-8), at increasing $NAD^+$ concentrations (0.5–100 µM). Lane 9 – control lane to show the positions of the substrate and 24-mer product in denaturing 20% PAG.

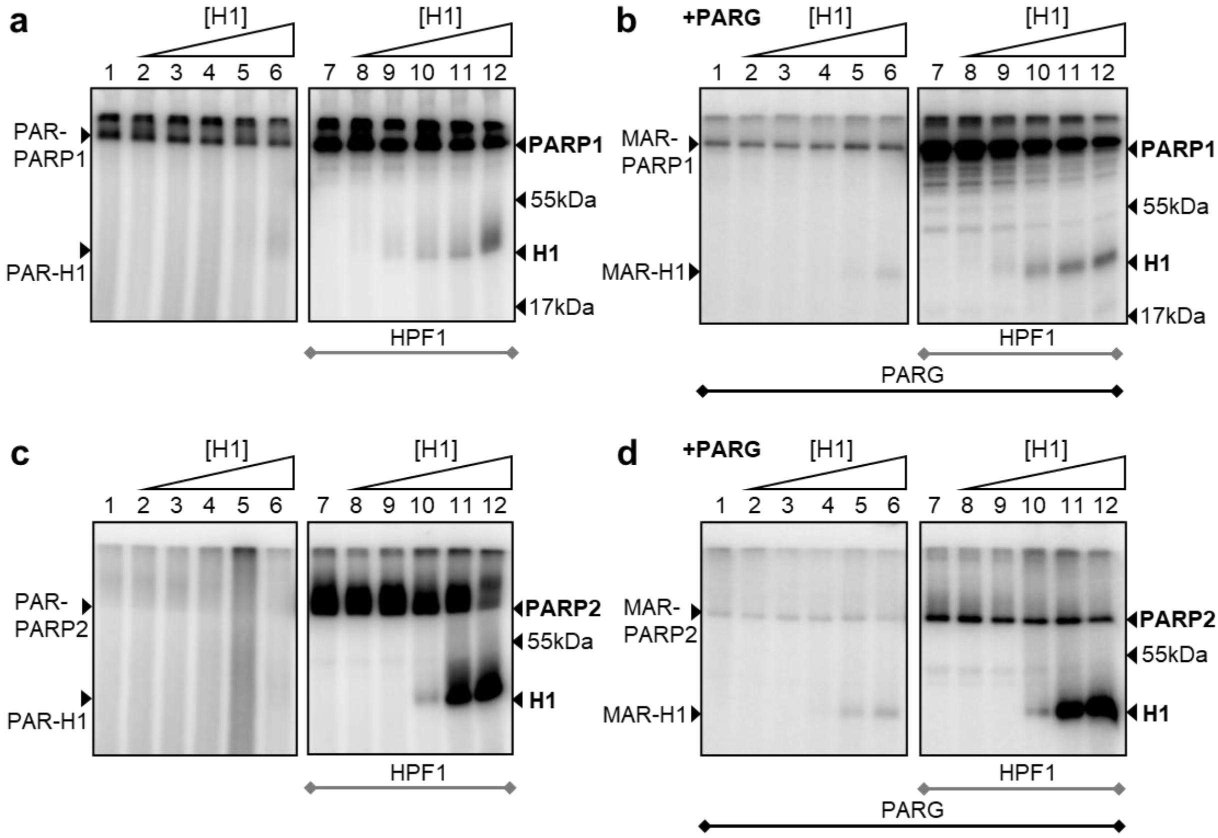


**Supplementary Fig. 8.** PARylation of histone H1 by PARP1 and PARP2 depends on its concentration and presence of HPF1.

(**a, b**) Autoradiograms show covalent attachment of $^{32}$P-labeled PAR (a) or MAR after treatment with 1 µM PARG (b) to protein targets after incubation of 100 nM PARP1 with 1 µM [$^{32}$P]NAD$^{+}$, 100 nM gap27-DNA227 and increasing concentrations of H1 (150 nM – 2.4 µM) in the absence (lanes 1–6) or presence (lanes 7–12) of 600 nM HPF1.

(**c, d**) The same experiments performed with 300 nM PARP2. Positions of the modified and unmodified forms of PARP1, PARP2 and H1 in 20% SDS-PAG, and of the molecular weight markers, are indicated to the left and right of the gel images. In the absence of HPF1, modification of H1 is detectable only at its highest concentrations, whereas in the presence of HPF1 it is efficient for both enzymes and particularly pronounced for PARP2.

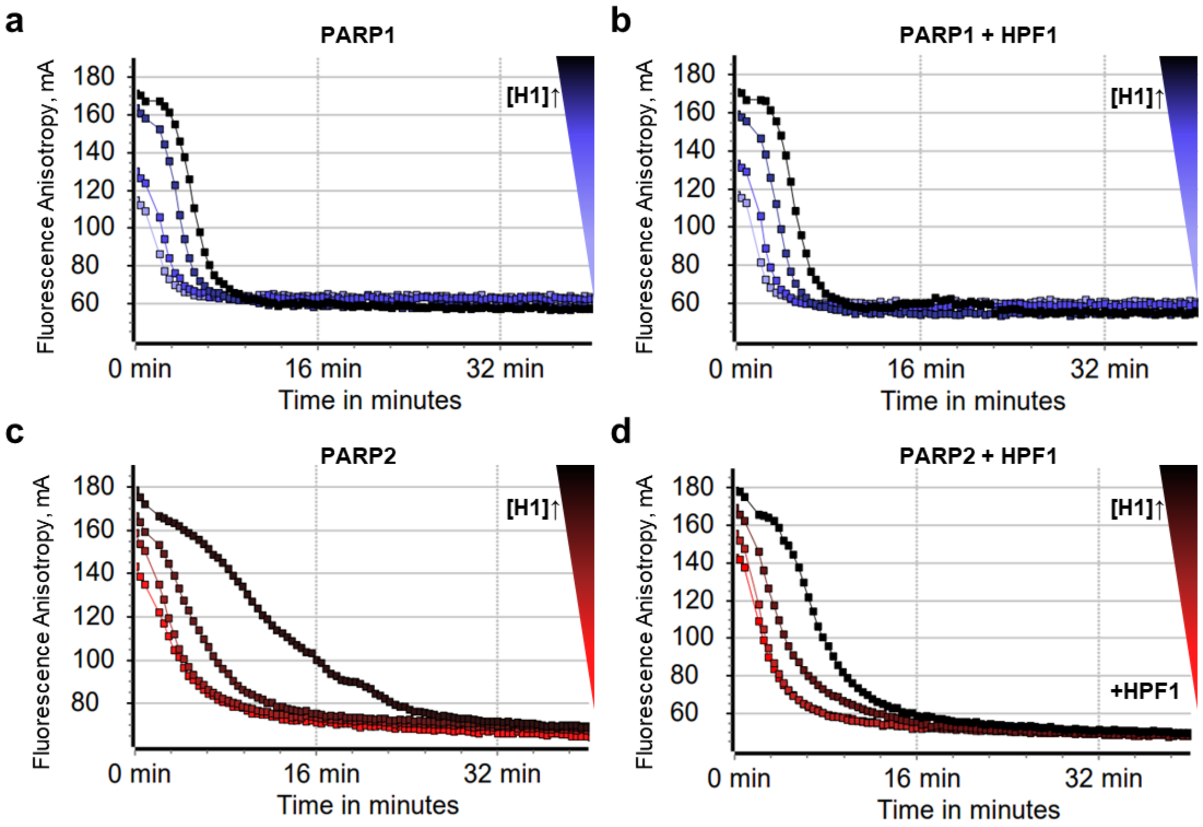


**Supplementary Fig. 9.** H1 slows the release of PARP1 and PARP2 from the nucleosomal substrate without changing the final signal.

Fluorescence anisotropy of FAM-labelled gap27-NCP227 (25 nM) prebound to 50 nM PARP1 (**a, b**) or 50 nM PARP2 (**c, d**) in the presence of 100 nM HPF1 and increasing concentrations of H1 (50–400 nM, color gradient), decreased after initiation of the PARylation reaction by $NAD^+$ addition. Increasing H1 concentrations slow down the dissociation of PARP1/PARP2 induced by automodification, while having no effect on the final signal (detected without H1).

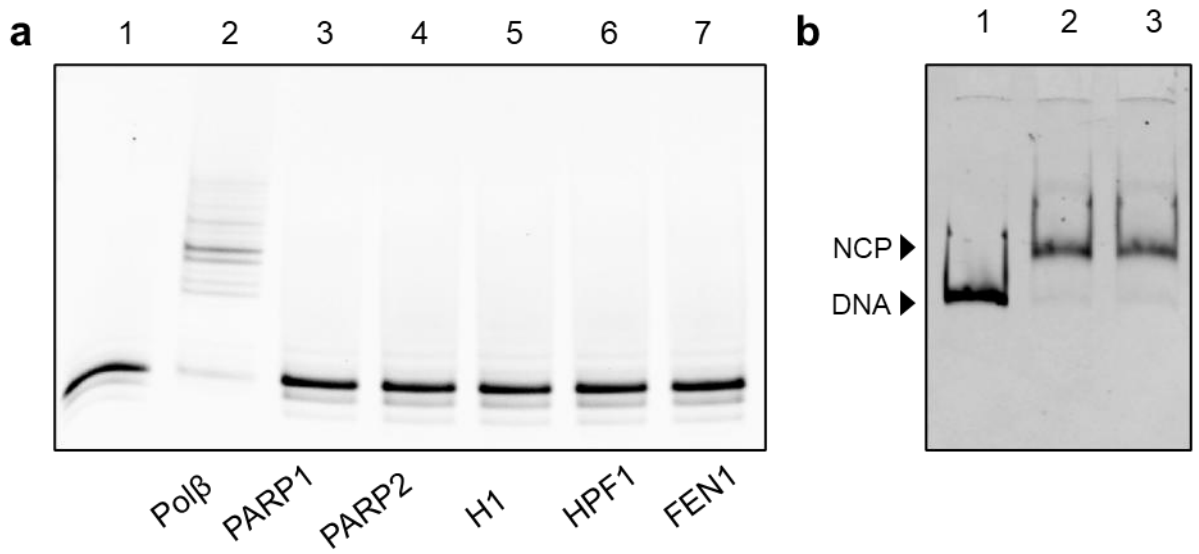


**Supplementary Fig. 10.** Control of the polymerase activity of the purified proteins and homogeneity of the reconstituted nucleosomes.

(**a**) gap27-DNA227 (50 nM) was incubated in the presence of four dNTPs (100 μM each) with 80 nM Polβ (lanes 1, 2) or 100 nM PARP1, 100 nM PARP2, 90 nM H1, 200 nM HPF1 or 200 nM FEN1 (lanes 3–7) as indicated below the gel. None of the accessory proteins displays detectable DNA polymerase activity.

(**b**) EMSA on a 4% native PAG shows the homogeneity of DNA227 (lane 1), NCP227 (lane 2) and nick-NCP227 (lane 3) preparations. The positions of the naked DNA and nucleosome are indicated to the left of the gel image.

### Supplementary Notes

**Supplementary Note 1 (related to Supplementary Figure 1):**
**Kinetic measurements of Polβ activity.**

The reaction of DNA synthesis via gap filling catalyzed by Polβ was carried out in reaction mixtures containing 200 nM DNA/NCP substrate, 4 nM Polβ and 500 nM dTTP. Reactions were initiated by addition of dTTPs and stopped after 20 s, 30 s, 40 s, 1 min, 3 min and 5 min of incubation by adding 5 µL aliquot of the reaction mixture to 5 µL of 8 M urea with 25 mM EDTA. To confirm single nucleotide incorporation, control reaction with ddTTP was used. Reaction products were separated by electrophoresis on 15% denaturing polyacrylamide gels. Gels were visualized using a Typhoon FLA 9500 scanner (GE Healthcare Life Sciences, USA) with excitation at 495 nm and emission at 520 nm. The relative intensities of bands corresponding to the substrate and products were quantified using Quantity One 4.6.6 software (Bio-Rad, USA) as follows: the total (raw) signal of the band of substrate or product (indicated for each product in the electropherograms) was quantified and the same-size background signal of gel in the respective lane was subtracted from the raw signal. All of the experiments were performed in three independent replicates. The data were plotted and fitted to exponential growth equation:
$[P] = A(1 - e^{-kt})$, where [P] is product concentration, A is maximum yield of product, k is observed reaction rate constant and t is time.

**Supplementary Note 2 (related to Supplementary Figure 4):**
**Electrophoretic mobility shift assay.**

To detect binding of Polβ to nucleosome, an electrophoretic mobility shift assay (EMSA) was used. The protein at increasing concentrations (50−200 nM) was incubated with FAM-labeled DNA227, gap27-DNA227, NCP227 or gap27-NCP227NCP227/gap27-NCP227 (50 nM) in a 10 µL mixture containing 50 mM Tris-HCl, pH 8.0, 50 mM NaCl, 5 mM $MgCl_2$ and 1 mM DTT at room temperature for 10 min. After the addition of Ficoll and bromophenol blue (to a final concentration of 5% and 0.1%, respectively), the incubation mixtures were electrophoresed at 4 °C on 5% non-denaturing PAG in a 30 mM Tris-Borate-EDTA buffer. Gels were imaged on a Typhoon FLA 9500 in the FAM channel.

**Supplementary Note 3 (related to Supplementary Figure 9):**
**Fluorescence studies of PARP1/PARP2 dissociation from complexes with DNA during PARylation.**

Fluorescence anisotropy measurements of FAM-labeled gap-NCP were performed in the absence and presence of PARP1, PARP2, HPF1 and increasing concentrations of H1. Briefly, a mixture containing 25 nM FAM-labeled gap27-NCP227, 100 nM HPF1 and 50 nM PARP1 or PARP2 in a buffer (50 mM NaCl, 50 mM Tris–HCl, pH 8.0, 5 mM $MgCl_2$ and 5 mM DTT) was prepared on ice in a 384-well plate and incubated at room temperature for 10 min. When indicated, samples were supplemented with histone H1 (50, 100, 200, 400 nM). The fluorescent probes were excited at 482 nm (482–16 filter plus dichroic filter LP504), and the fluorescence intensities were detected at 530 nm (530–40 filter) to measure the fluorescence anisotropy of FAM. To initiate the PARylation reacton, $NAD^+$ was added to each of samples to final concentration of 100 µM. Each measurement consisted of 50 flashes per well, and the resulting values of fluorescence anisotropy were automatically averaged. The measurements in each well were done with intervals of 20 sec during 40 min.
